# Epithelial FOXP3 Drives Pancreatic Fibrosis through O-glycosylated IL-6

**DOI:** 10.1101/2025.11.19.689208

**Authors:** Ruining Gong, Minghan Ren, Junjin Wang, Wenqing Zhang, Shuai Ping, Ke Lei, Qian Yu, Leina Ma, Chao Gao, Chenyang Zhao, Shunli Fu, Jing Xue, Jigang Wang, Hao Sun, Jinsuo Chen, Shugang Zhang, Mingyang Liu, Tiansuo Zhao, Ashok Saluja, Zhan Yang, He Ren

## Abstract

**BACKGROUND & AIMS:** The transcription factor FOXP3, known for specifying regulatory T cell fate, is unexpectedly expressed in the epithelium of precancerous pancreatic lesions. Here we investigate its non-immune role in initiating pancreatic fibrosis, a process driving therapeutic resistance and organ failure in pancreatic ductal adenocarcinoma (PDAC) with unknown origins in early neoplasia.

**METHODS:** Using human tissues and genetically engineered mouse models, we analysed FOXP3 expression in premalignant lesions. We employed epithelial-specific FOXP3 knockout and knock-in strategies to determine its functional impact on fibrogenesis and neoplasia progression. Mechanistic studies included chromatin immunoprecipitation, glycomic analyses, and signalling assays.

**RESULTS:** FOXP3 was consistently expressed in the epithelial compartment of human and murine precancerous pancreas. Epithelial-specific deletion of FOXP3 attenuated pancreatic fibrosis and delayed neoplasia, whereas its knock-in induced spontaneous stromal activation and accelerated PanIN progression. Mechanistically, epithelial FOXP3 directly transactivated the glycosyltransferase *GALNT1*. GALNT1, in turn, mediated O-glycosylation of interleukin (IL)-6, which was essential for its rapid secretion.

**CONCLUSIONS:** Our study establishes epithelial-derived FOXP3 as a master regulator of early pancreatic fibrocarcinogenesis. It drives a glycosylation-dependent amplification loop for IL-6 signalling, orchestrating sustained stromal activation. This pathway represents a promising target for intercepting pancreatic fibrosis and carcinogenesis at its origin.

## INTRODUCTION

Forkhead box protein 3 (FOXP3) is unequivocally established as the master transcriptional regulator of regulatory T cells (Tregs)^1^, a role so fundamental that it was underscored by the 2025 Nobel Prize in Physiology or Medicine. This canonical identity, however, has largely confined our understanding of FOXP3 to the realm of immune tolerance^2^. Beyond the immune system, emerging evidence suggests that FOXP3 expression exhibits remarkable context-dependency across different cell types^3^. The functional consequences of this expression outside the immune compartment, particularly in epithelial cells during early pathogenesis, remain largely underexplored.

Pancreatic fibrosis is a pathological hallmark of both chronic pancreatitis and pancreatic ductal adenocarcinoma (PDAC), contributing to organ dysfunction, therapeutic resistance, and poor patient outcomes^4,5^. Although activated pancreatic stellate cells (PSCs) and fibroblasts are well-established as the primary effectors producing extracellular matrix, the upstream signals that trigger their transition from quiescent to profibrotic states remain poorly defined^6^. While inflammatory mediators derived from immune cells dominate the fibrotic landscape in late-stage PDAC^7,8^, early-stage lesions often develop without overt clinical symptoms—such as abdominal pain—suggesting that inflammation-independent mechanisms may underlie the initiation of fibrosis. Emerging evidence implicates epithelial cells as potential orchestrators of stromal remodelling^4,9^; however, their specific role in the onset of fibrosis remains largely underexplored.

In the course of investigating Tregs, we observed FOXP3 expression in epithelial cells during the early stages of pancreatic neoplasia, which we refer to as epithelial-FOXP3 (hereafter E-FOXP3). Since E-FOXP3^+^ epithelial cells are rare and spatially restricted within early lesions, they are particularly challenging to capture and characterize with conventional methodologies such as scRNA-seq, which often overlooks such rare but functionally significant cell populations^10–12^. Utilising genetically engineered mouse models (GEMMs) with pancreatic epithelial-specific FOXP3 knockout and knock-in, we demonstrate that epithelial FOXP3 deletion attenuates stress-induced fibrosis, while its overexpression is sufficient to initiate fibrotic programs and accelerate precancerous progression. Notably, E-FOXP3 drives pro-fibrotic signaling, functionally opposing its classical immunosuppressive role in Tregs^13^.

Mechanistically, E-FOXP3 directly transactivates the glycosyltransferase polypeptide *N*-acetylgalactosaminyltransferase 1 (GALNT1), driving glycosylation of IL-6 and enabling its rapid secretion. This glycosylated IL-6 (G-IL-6) directly activates PSCs to drive inflammatory fibrosis during early pancreatic preneoplasia. Thus, we redefine E-FOXP3^+^ epithelial cells as proto-fibrogenic ‘trigger cells’, unveiling a non-canonical, glycosylation-dependent pathway that bridges epithelial dysfunction to stromal activation in early pancreatic neoplasia.

## RESULTS

### E-FOXP3 marks preneoplastic lesions and promotes fibrotic progression in pancreatic tumorigenesis

Although FOXP3 expression has been previously documented in pancreatic ductal adenocarcinoma (PDAC) cells, its epithelial-specific role within preneoplastic lesions remains poorly understood ^14–16^. To address this knowledge gap, we first evaluated FOXP3 expression in human pancreatic tissues. We identified FOXP3 expressed in epithelial cells (FOXP3⁺CK19⁺) within pancreatic intraepithelial neoplasia (PanIN) lesions from human PDAC samples (Fig. 1A, B). Although this epithelial subpopulation was relatively rare, the proportion of FOXP3⁺ epithelial cells among total epithelial cells increased significantly with lesion progression (Fig. 1C). To investigate whether E-FOXP3 accumulates in precancerous lesions during natural disease course, we further analyzed pancreatic tissues from human donors presenting with chronic inflammation, PanIN lesions, and overt stromal fibrosis. Immunohistochemical (IHC) analysis revealed that E-FOXP3 was expressed in PanIN epithelial cells but was absent in normal acinar or ductal epithelia (Supplementary Fig. S1A, B), indicating a lesion-specific expression pattern.

**Figure 1.**
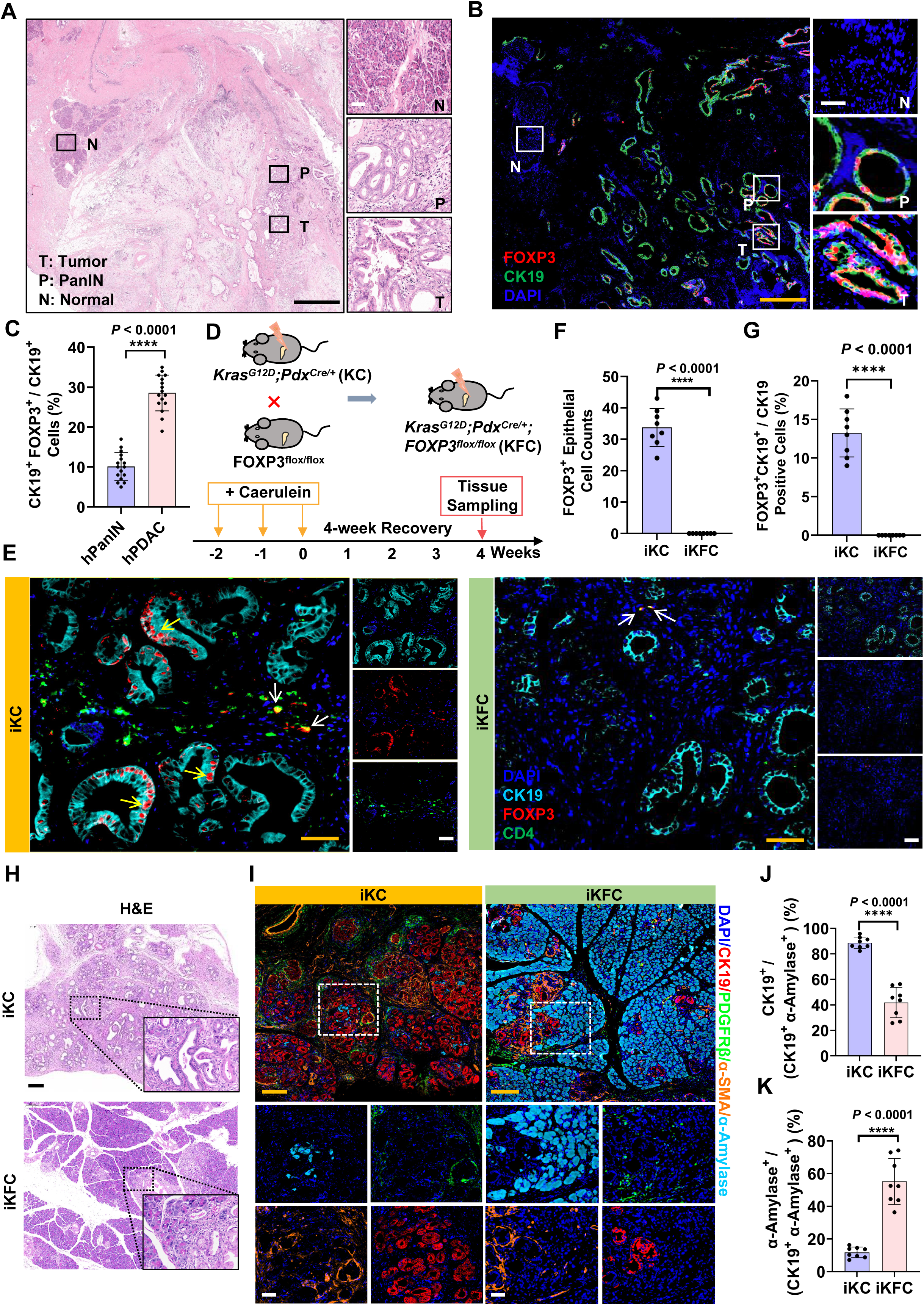
FOXP3 expression in human and mouse premalignant pancreatic lesions. **A,** Representative high-magnification images of haematoxylin and eosin (H&E) staining of human PDAC tissues, showing sections of normal (N), PanIN (P), and tumour (T). Scale bars: 250 μm (black), 50 μm (white). **B**, High-magnification multiplex immunofluorescence images of human PDAC tissues showing co-localization of CK19 (green), FOXP3 (red), DAPI (blue) of N, P, and T regions. Scale bars: 250 μm (orange), 50 μm (white). **C,** Percentage of FOXP3^+^CK19⁺ cells relative to total CK19⁺ cells in the pancreases of human (*n* = 16). **D,** Schematic diagram of the caerulein-induced chronic pancreatitis model in KC and KFC mice. KC, *LSL-Kras^G12D^*; *Pdx1-Cre* mice; KFC, *Kras^G12D^; FOXP3^flox/flox^; Pdx1-Cre* mice. **E**, Multiplex immunofluorescence (mIF) staining for FOXP3 (red), CD4 (green), and CK19 (cyan) in pancreatic sections from iKC and iKFC mice. Scale bar: 50 μm (orange), 50 μm (white). (yellow arrow: FOXP3^+^ epithelial cells; white arrow: CD4⁺ regulatory T cells). **F**, Bar graphs showing the total FOXP3⁺ epithelial cells in the pancreases of mice (*n* = 8). **G**, Percentage of FOXP3^+^CK19⁺ cells relative to total CK19⁺ cells in the pancreases of mice (*n* = 8). **H**, H&E staining of caerulein-induced chronic pancreatitis in KC and KFC mice. Scale bar: 50 μm. **I**, mIF staining for α-amylase (cyan), PDGFRβ (green), α-SMA (orange), and CK19 (red) in pancreatic sections from KC and KFC mice. Scale bars: 200 μm (orange), 50 μm (white). **J,** Bar graphs showing the percentage of α-SMA⁺ cells relative to PDGFRβ⁺ cells in the pancreases of mice (*n* = 8). **K,** Percentage of CK19⁺ cells relative to total CK19⁺ and α-amylase⁺ cells in the pancreases of mice (*n* = 8). Data are presented as mean ± SD; n = 3 biological replicates. \*\*\**P* < 0.001, \*\*\*\**P* < 0.0001; paired two-tailed t-test.

To model the development of precancerous lesions, we employed the KC mouse model (*LSL-Kras^G12D^; Pdx1-Cre*), a well-established genetic model of early PDAC (Supplementary Fig. S1C). We performed IHC on pancreatic tissues from healthy wild-type (WT) mice, KC mice, WT mice subjected to caerulein-induced chronic pancreatitis (WT-CP), and KC mice with caerulein-induced chronic pancreatitis (KC-CP). FOXP3 expression was undetectable in WT pancreatic tissues (Supplementary Fig. S1D). In contrast, cytoplasmic FOXP3 expression was observed in both acinar and ductal cells of WT-CP mice (Supplementary Fig. S1E), a pattern also recapitulated in KC mice (Supplementary Fig. S1F). Notably, nuclear FOXP3 expression in ductal epithelial cells was exclusively detected in mice harboring both oncogenic *Kras* mutation and caerulein-induced inflammation (KC-CP), mirroring the low-frequency expression observed in human PanIN samples (Supplementary Fig. S1G-1I). These findings establish E-FOXP3 as a marker selectively induced in preneoplastic lesions.

Given the association between E-FOXP3 and early pancreatic lesions, we generated a pancreas-specific conditional knockout mouse model (*Kras^G12D^; FOXP3^flox/flox^; Pdx1-Cre*, hereafter denoted KFC) to investigate the functional contribution of E-FOXP3 to fibrogenesis (Supplementary Fig. S2A). Chronic inflammation was induced in both KC and KFC mice via intraperitoneal caerulein administration (Fig. 1D). Notably, KFC mice exhibited significantly attenuated pancreatic inflammation and fibrosis compared to KC littermates, as evidenced by reduced weight loss, higher pancreas-to-body weight ratios, and softer tissue consistency (Supplementary Fig. S2B–D). Multiplex immunofluorescence (mIF) staining for FOXP3, cytokeratin 19 (CK19, epithelial cell marker), and CD4 (T cell marker) confirmed a substantial increase in E-FOXP3⁺ cells within PanIN lesions of KC mice, which was abrogated in KFC mice (Fig. 1E, F). Importantly, FOXP3 expression remained intact in CD4⁺ regulatory T cells (Tregs) in both KC and KFC mice (Fig. 1E), validating the epithelial-specific nature of the knockout. Histomorphological analysis via haematoxylin and eosin (H&E) staining revealed that KFC pancreatic tissues retained a greater proportion of normal acinar architecture (Fig. 1H and Supplementary Fig. S2E). Consistent with this, histopathological evaluations—including Alcian blue, Sirius red, and Masson’s trichrome staining—demonstrated that epithelial-specific deletion of FOXP3 significantly mitigated pancreatic fibrotic progression (Supplementary Fig. S2F-K). Whereas KC mice developed extensive high-grade PanIN lesions accompanied by substantial acinar replacement with dense fibrotic stroma, KFC pancreata exhibited only focal low-grade PanINs with markedly reduced stromal expansion (Supplementary Fig. S2H-K).

Further multiplex immunofluorescence staining for CK19, α-amylase, platelet-derived growth factor receptor β (PDGFR-β), and α-smooth muscle actin (α-SMA) revealed that KFC mice maintained a higher abundance of α-amylase⁺ acinar cells and showed reduced density of α-SMA⁺PDGFR-β⁺ activated fibroblasts relative to KC mice (Fig. 1I-K and Supplementary Fig. S2L). These results indicate that loss of E-FOXP3 preserves epithelial integrity and suppresses stromal activation in the pancreas.

### FOXP3^+^ epithelial cells are sufficient to drive pancreatic fibrosis

To determine whether E-FOXP3 expression alone is sufficient to initiate fibrotic remodelling, we generated a pancreatic epithelial cell–specific FOXP3 knock-in mouse model (*FOXP3^R26^; Pdx1-Cre*, hereafter referred to as RC) using CRISPR/Cas9 technology (Fig. 2A). Gross morphological analysis revealed no overt differences in pancreatic appearance between *FOXP3^R26^* (R) and RC mice (Supplementary Fig. S3A). However, starting at 10 weeks of age, RC mice displayed significant weight loss compared to their R littermates (Supplementary Fig. S3B), along with a reduced pancreas-to-body weight ratio indicative of pancreatic atrophy (Supplementary Fig. S3C), and signs of reduced vitality despite the absence of visible pancreatic mass lesions (Supplementary Fig. S3D). IHC analysis confirmed robust FOXP3 expression in pancreatic epithelial cells of RC mice, consistent with their genotype (Fig. 2B and 2C). Histopathological evaluation revealed spontaneous emergence of focal PanIN lesions in RC mice, occurring without any external inflammatory stimulus. These lesions were accompanied by extensive stromal fibrosis–a phenotype absent in R controls (Fig. 2D and 2E). Consistent with prior observations, the abundance of E-FOXP3^+^ epithelial cells increased with pathological progression from PanIN1 to PanIN3 lesions (Supplementary Fig. S3E, F). Further histological assessment demonstrated that RC mice developed marked pancreatic inflammation, acinar-to-ductal metaplasia (ADM), and a higher frequency of PanIN lesions compared to R mice (Fig. 2F-2G). Alcian Blue, Masson’s trichrome, and Sirius Red staining confirmed substantial fibrotic stromal infiltration in the RC pancreata, while R mice retained normal pancreatic architecture (Fig. 2H-2M). Multiplex immunofluorescence staining revealed that the fibrotic stroma in RC mice comprised activated fibroblasts, consistent with the profile observed in *Kras*-driven models (Fig. 2N-2P).

**Figure 2.**
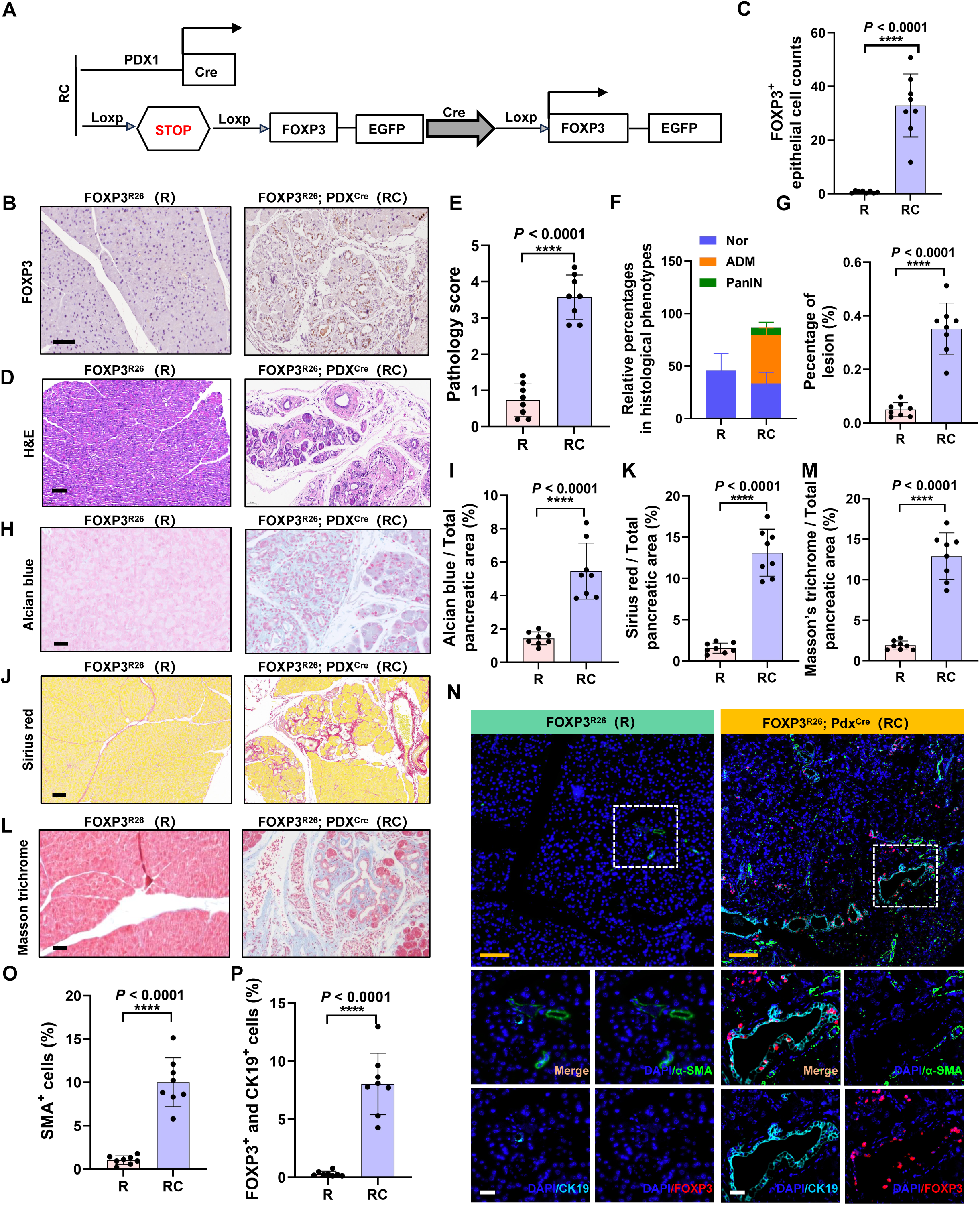
FOXP3^+^ epithelial cells sufficiently induce pancreatic fibrosis. **A,** Schematic diagram of the RC conditional transgenic mouse model. **B and C,** IHC staining for FOXP3 (B) and corresponding quantification (C) of pancreatic tissues from R (control) and transgenic (RC) mice. Scale bar: 100 μm. RC, *FOXP3^R26^; Pdx1-Cre* mice; R, *FOXP3^R26^* control mice. **D,** H&E staining of pancreatic tissues from R and RC mice. Scale bar: 50 μm. **E-G,** Quantification of pancreatic pathology in R and RC mice (*n* = 8): pathology scores (E), relative percentages of histological phenotypes (F), and percentage of fibrotic lesions (G). **H,** Alcian blue staining of pancreatic tissues from R and RC mice. Scale bar: 50 μm. **I,** Bar graphs showing the percentage of Alcian blue–positive areas in the pancreases of R and RC mice. **J,** Sirius red staining of pancreatic tissues from R and RC mice. Scale bar: 50 μm. **K,** Bar graphs showing the percentage of Sirius red-positive staining in the pancreases of R and RC mice. **L,** Masson’s trichrome staining of pancreatic tissues from R and RC mice. Scale bar: 50 μm. **M**, Bar graphs showing the percentage of Masson’s trichrome–positive lesions in the pancreases of R and RC mice. **N-P,** Representative mIF images and quantitative analysis of CK19, FOXP3, and α-SMA in pancreatic tissues from R and RC mice (*n* = 8). Scale bars: 200 μm (orange), 50 μm (white). Data in (C), (E), (F), (G), (I), (K), (M), (O) and (P) are represented as mean ± SD for n = 3 biological replicates. \*\*\*\**P* < 0.0001, determined by a paired two-tailed t test.

### E-FOXP3 remodels the fibroblast landscape and promotes an inflammatory microenvironment

To elucidate the cellular mechanisms underlying E-FOXP3-driven fibrogenesis, we performed single-cell RNA sequencing (scRNA-seq) on pancreata from *KC* mice (Fig. 3A). Clustering and dimensionality reduction via t-distributed stochastic neighbor embedding (t-SNE) resolved 15 transcriptionally distinct cell populations (Fig. 3B, C; Supplementary Fig. S3A), including inflammatory cancer-associated fibroblasts (iCAFs; marked by *Svep1*), myofibroblastic CAFs (myCAFs; *Dkk2, Pdgfrl*), antigen-presenting CAFs (apCAFs; *Fras1, Upk1b, Lrp2, Gm32587*), pancreatic stellate cells (PSCs; *Bpgm, Sparcl1, Dlc1*), acinar cells (*Prss2, Cpa1*), ductal cells (*Slc28a3, Mmp7, Dcdc2a, Cldn4*), endocrine cells (*Neurod1*), endothelial cells (*Cdh5, Klhl4, Flt4*), dendritic cells (DCs; *Gm36723, Gm35853, Gcsam, Gpr4*), monocytes (*Il1r2*), plasma cells (*Gm5547, Sec11c, Tnfrsf17, Oosp1, Nuggc, Mzb1*), M1 macrophages (*Cd80, Htr7*), M2 macrophages (*Cd163, Gm30489, Aoah*), T regulatory cells (Tregs; *Cd5, Skap1, Bcl11b*), and B cells (*Tmem163, Fcer2a, H2-DMb2*).

**Figure 3.**
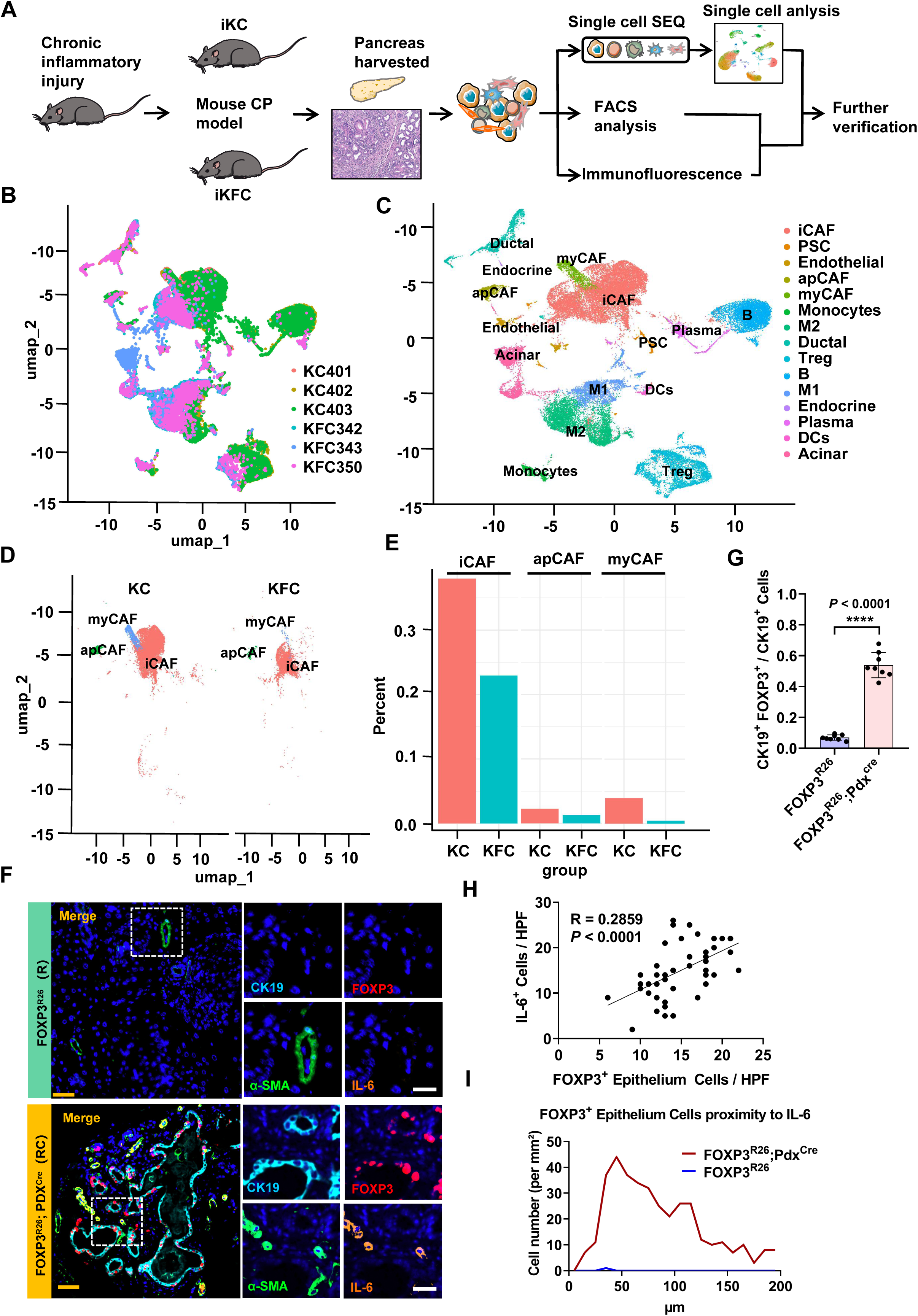
Single-cell transcriptomic analysis of chronic pancreatitis tissue in KC and KFC mice. **A,** Schematic diagram illustrating tissue collection and the single-cell transcriptomic analysis workflow for chronic pancreatitis. **B and C,** Uniform Manifold Approximation and Projection (UMAP) plot of 40,394 pancreatic cells, coloured by individual sample groups (B) and by annotated cell types (C). **D,** UMAP plot showing the distribution of iCAF, myCAF, and apCAF subsets in the KC and KFC groups. **E,** Bar graph comparing the proportions of iCAF, myCAF, and apCAF subsets between KC and KFC mice, with the most significant difference observed in iCAF. **F**, mIF staining showing co-localisation of CK19^+^FOXP3^+^ cells and IL-6^+^α-SMA^+^ fibroblasts in pancreatic tissues of R and RC mice. Scale bars: 100 μm (orange), 50 μm (white). **G**, Proportion of CK19^+^FOXP3^+^ cells among CK19^+^ cells in the pancreas of R and RC mice. **H**, Correlation between FOXP3^+^ and IL-6^+^ cells in the RC mouse pancreas. **I,** Distance distribution between FOXP3^+^ and IL-6^+^ cells in the pancreas of R and RC mice.

Comparative analysis of fibroblast subpopulations between KC and KFC models revealed substantial shifts in subtype composition, most notably within iCAFs. Uniform Manifold Approximation and Projection (UMAP) visualization highlighted distinct clustering patterns of iCAFs, myCAFs, and apCAFs based on epithelial FOXP3 status (Fig. 3D). Quantitative evaluation confirmed significant enrichment of iCAFs in KC compared to KFC mice (Fig. 3E), indicating that E-FOXP3 promotes the expansion of this inflammatory fibroblast subset. In contrast, macrophage populations—often associated with fibrosis in other settings^17^—were reduced in KC pancreata (data not shown), suggesting that macrophage infiltration may play a limited role in early fibrocarcinogenesis and likely contributes at later stages. These findings implicate iCAFs as the dominant fibroblast subtype contributing to stromal fibrosis in KC mice and suggest that epithelial FOXP3 expression shapes the fibrotic microenvironment by sustaining this specific fibroblast population.

Given the unexpected reduction in classical immune mediators and the prominence of iCAFs, we hypothesized that epithelial FOXP3 may drive fibrogenesis through direct modulation of fibroblast activity rather than through canonical immune pathways. To test this hypothesis and establish the sufficiency of epithelial FOXP3 in initiating these changes, we turned to a gain-of-function model. We performed multiplex immunofluorescence imaging on pancreatic tissues from *FOXP3^R26^; Pdx1-Cre* (RC) mice, which enable targeted FOXP3 expression in pancreatic epithelial cells. We observed a significant increase in CK19^+^FOXP3^+^ epithelial cells in RC compared to R mice, with these cells predominantly localized within high-grade PanIN lesions (Fig. 3F, G). RC pancreata also exhibited elevated densities of IL-6^+^ cells, and IL-6^+^ cell abundance correlated positively with CK19^+^FOXP3^+^ epithelial cell counts (Fig. 3H). We frequently detected IL-6^high^α-SMA^low^ cells—a signature of inflammatory cancer-associated fibroblasts (iCAFs)^18^—in close proximity to FOXP3^+^ epithelial cells, suggesting their involvement in the immune and fibrotic responses characteristic of RC pancreata. Spatial analysis confirmed enhanced clustering of IL-6^+^ cells adjacent to CK19^+^FOXP3^+^ epithelial structures (Fig. 3I; Supplementary Fig. S4B). Together, these data reveal that E-FOXP3 expression drives the expansion of a specific inflammatory fibroblast subset (iCAFs) and promotes a localized IL-6-enriched microenvironment, thereby facilitating stromal fibrogenesis and PanIN progression.

### E-FOXP3–IL-6 signaling drives inflammatory fibroblast activation

To determine whether E-FOXP3 promotes an inflammatory phenotype in pancreatic stellate cells (PSCs) via IL-6, we co-cultured FOXP3-overexpressing human pancreatic ductal epithelial cells (hTERT-HPNE) with PSCs (Fig. 4A). Co-culture induced pronounced PSC activation, characterized by elevated IL-6 expression and reduced levels of α-SMA and PDGFRα, consistent with a shift toward an inflammatory CAF (iCAF) phenotype (Fig. 4B, Supplementary Fig. S5A). These changes were corroborated at both transcriptional and protein levels (Fig. 4C, D).

**Figure 4.**
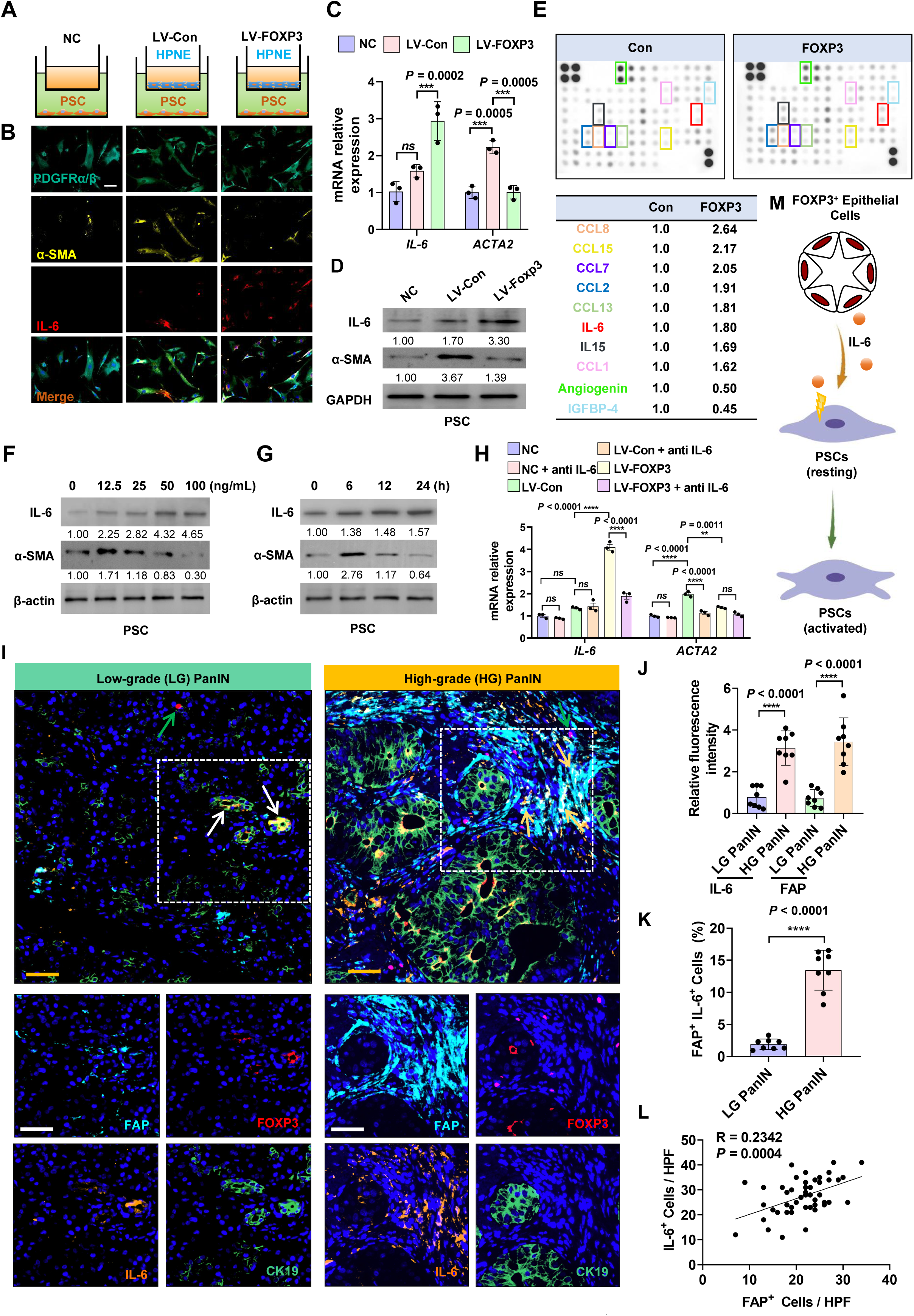
Overexpression of FOXP3 in pancreatic epithelial cells activates fibroblasts by enhancing IL-6 secretion. **A,** Schematic of the co-culture strategy between epithelial cells and PSCs. **B,** Immunofluorescence staining of PSC activation markers after 24 h of co-culture. Scale bar: 100 μm. **C and D,** Relative mRNA and protein expression levels of IL-6 and ACTA2 (α-SMA), markers of PSC activation, after 24 h of co-culture. **E,** Cytokine array showing the levels of secreted cytokines in the supernatants of hTERT-HPNE cells with varying FOXP3 expression, along with quantitative analysis. **F,** Protein expression levels of PSC activation markers following stimulation with different concentrations of IL-6 (100 ng/mL). **G,** Protein expression levels of PSC activation markers after treatment with 100 ng/mL IL-6 at various time points. **H,** Relative mRNA levels of PSC activation markers after co-culture with hTERT-HPNE cells expressing different levels of FOXP3 with or without IL-6 neutralising antibody treatment. **I**, mIF staining showing colocalisation of IL-6^+^FOXP3^+^, CK19^+^FOXP3^+^ epithelial cells and IL-6^+^FAP^+^ fibroblasts in human low grade (LG) and high grade (HG) PanIN tissues. Scale bars: 100 μm (orange), 50 μm (white). White arrow: FOXP3^+^ epithelial cells; green arrow: Extra-ductal FOXP3⁺ cells; yellow arrow: FAP^+^IL-6^+^ cell. **J**, Relative fluorescence Intensity of IL-6 and FAP expression level in LG and HG PanIN lesions. **K**, Proportion of FAP^+^IL-6^+^ cells in human LG and HG PanIN tissues. **L**, Correlation between FAP^+^ and IL-6^+^ cells in human tissues. **M,** Schematic diagram showing the proposed mechanism by which FOXP3^+^ pancreatic epithelial cells activate PSCs via IL-6 secretion. Data in (C) are represented by a paired two-tailed *t* test. (H) are represented by one-way ANOVA with Tukey’s post-hoc test. \*\**P* < 0.01, \*\*\**P* < 0.001, \*\*\*\**P* < 0.0001 vs. corresponding control.

To identify soluble mediators underlying this activation, we profiled conditioned media from FOXP3⁺ hTERT-HPNE cells using protein microarrays. We observed marked upregulation of inflammatory cytokines including IL-6, CCL2, CCL8, and CCL15, alongside downregulation of angiogenin and IGFBP-4 (Fig. 4E). ELISA further validated these secretory alterations (Supplementary Fig. S5B). To pinpoint which cytokine mediates E-FOXP3–driven inflammatory activation in PSCs, we treated PSCs individually with each candidate factor. Only IL-6 stimulation simultaneously enhanced IL-6 expression and suppressed α-SMA in PSCs (Supplementary Fig. S5C), suggesting IL-6 as a key mediator of FOXP3-induced PSC activation.

Given the established role of IL-6 as a core iCAF marker and mediator of pancreatic tumor progression^19,20^, we treated PSCs with recombinant IL-6 and assessed expression of iCAF markers. IL-6 mRNA increased dose-dependently, while ACTA2 (encoding α-SMA) showed transient upregulation at low concentrations but declined at higher levels (Supplementary Fig. S5D). Time-course analysis under fixed IL-6 concentration (100 ng/mL) revealed sustained induction of IL-6 but a late decrease in ACTA2 (Supplementary Fig. S5E), suggesting progressive commitment to an iCAF state. Western blot analysis confirmed dynamic changes in protein expression (Fig. 4F, G). Critically, IL-6 neutralization with a specific antibody during co-culture abrogated PSC activation (Fig. 4H), establishing IL-6 as the central mediator of this phenotypic shift.

To further investigate whether FOXP3⁺ epithelial cells promote PSC activation and initiate an IL-6 feedback loop through paracrine signaling, we performed multiplex immunofluorescence staining on pancreatic tissues. During the low-grade PanIN stage, IL-6 was predominantly localized within FOXP3⁺ epithelial cells (white arrows) and was largely absent in extra-ductal FOXP3⁺ cells (green arrows). In contrast, at the high-grade PanIN stage, IL-6 was mainly detected in FAP⁺ cancer-associated fibroblasts (yellow arrows) and was scarcely present in ductal epithelia (Fig. 4I). Both IL-6 and FAP expression were significantly elevated in high-grade PanIN lesions compared to earlier stages (Fig. 4J). Moreover, the abundance of FAP⁺IL-6⁺ cells was markedly increased in high-grade PanIN relative to low-grade PanIN (Fig. 4K). A positive correlation was observed between IL-6⁺ cell density and FAP⁺ epithelial cell counts (Fig. 4L), suggesting that early FOXP3⁺ epithelial cells may initiate PSC activation via IL-6 secretion, and activated PSCs in turn amplify IL-6 production to propagate fibrotic remodeling (Fig. 4M).

### E-FOXP3 transactivates GALNT1 to drive IL-6 glycosylation and secretion

To investigate whether E-FOXP3 enhances IL-6 secretion through transcriptional regulation, we first assessed IL-6 mRNA expression in hTERT-HPNE cells. While E-FOXP3 overexpression significantly upregulated the known target *p21*^21^, it did not alter IL-6 transcript levels (Supplementary Fig. S6A). Chromatin immunoprecipitation (ChIP)-PCR further confirmed that E-FOXP3 does not bind the -2000 bp region of IL-6 promoter but does occupy the p21 promoter region (Supplementary Fig. S6B). Moreover, RNA stability assays indicated that E-FOXP3 does not affect IL-6 mRNA half-life (Supplementary Fig. S6C).

We next asked whether E-FOXP3 influences IL-6 secretion. Treatment with Brefeldin A, an inhibitor of protein secretion, reduced extracellular IL-6 and increased its intracellular accumulation. E-FOXP3 overexpression enhanced IL-6 secretion into the supernatant and decreased intracellular retention—effects that were abolished upon Brefeldin A treatment (Fig. 5A, B; Supplementary Fig. S6D). These data suggest that E-FOXP3 promotes IL-6 secretion without affecting its synthesis or stability.

**Figure 5.**
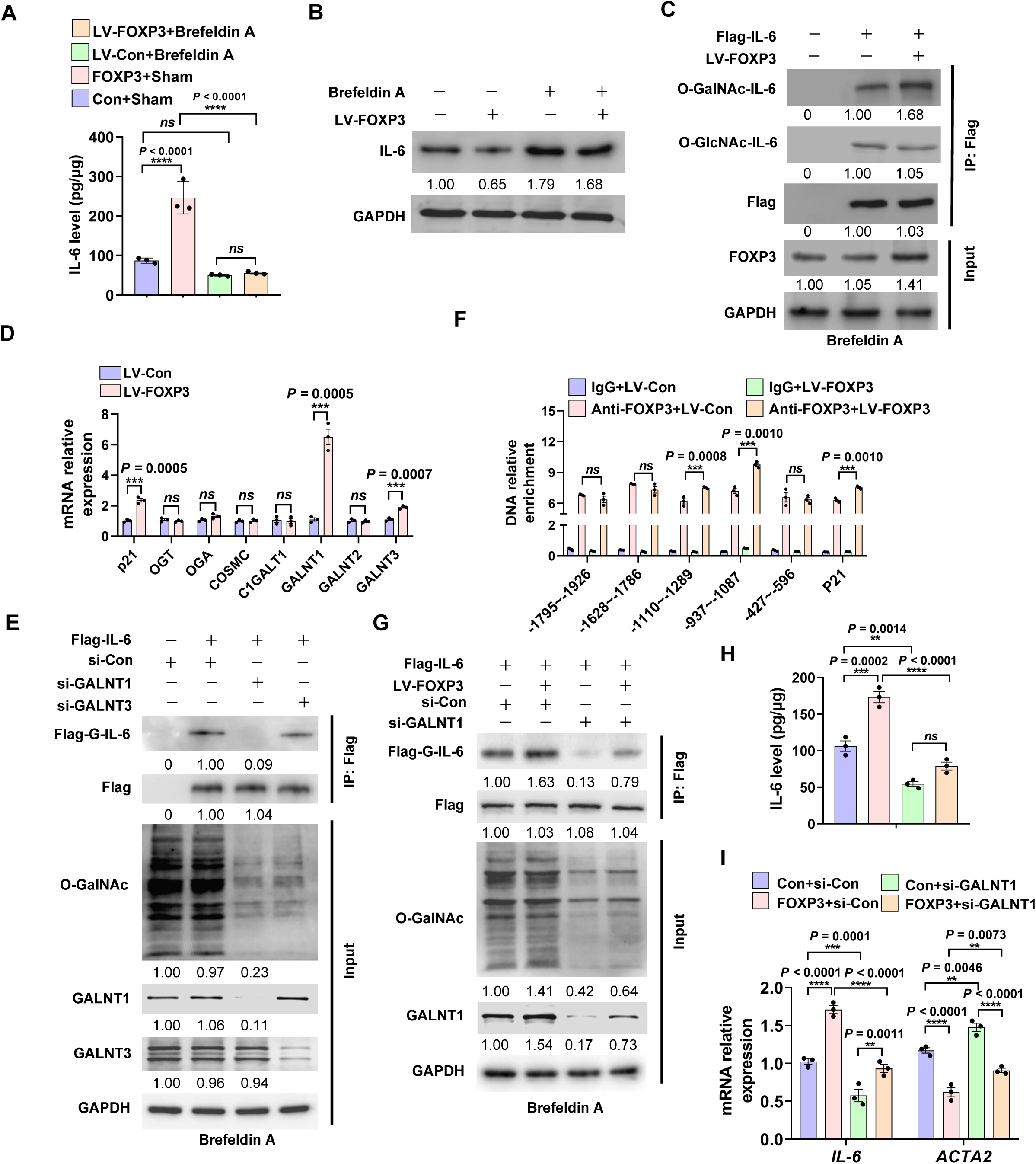
FOXP3 promotes the glycosylation modification and secretion of IL-6 by promoting GALNT1 transcription. **A and B,** hTERT-HPNE cells were infected with lentivirus-FOXP3 (LV-FOXP3) or control lentivirus (LV-Con) and treated with or without the protein transport inhibitor Brefeldin A for 24 h. IL-6 levels were measured in culture supernatants (A) and within hTERT-HPNE cells (B). **C,** hTERT-HPNE cells were transfected with the FLAG-tagged IL-6 vector and LV-FOXP3 lentivirus, followed by Brefeldin A treatment. Co-IP was performed to detect O-GalNAc-IL-6 and O-GlcNAc-IL-6. **D,** mRNA expression levels of O-glycosylation-related genes in LV-control and LV-FOXP3-infected hTERT-HPNE cells. **E,** hTERT-HPNE cells were transfected with the Flag-IL-6 vectors, GALNT1 and GALNT3 siRNAs, treated with Brefeldin. Cell lysate subjected to Co-IP with Flag antibody and immunoblotting was used to detect G-IL-6 and indicated proteins. **F,** ChIP assay of FOXP3 binding to the -2000 bp region of *GALNT1* promoter in hTERT-HPNE cells. **G,** hTERT-HPNE cells expressing Flag-IL-6 were transfected with LV-FOXP3 and si-GALNT1, Co-IP with Flag antibody and immunoblotting were used to detect G-IL-6 and indicated proteins. **H,** hTERT-HPNE cells were treated as in (g) and IL-6 levels in the culture supernatants were measured by ELISA. **I,** hTERT-HPNE cells transfected as in (g) were co-cultured with PSCs for 48 h. mRNA levels of *IL-6* and *ACTA2* (α-SMA) in PSCs were quantified by RT-qPCR. Data in (A), (D) and (F) are represented by a paired two-tailed *t* test. (H) and (I) are represented by one-way ANOVA with Tukey ’ s post-hoc test. \*\**P* < 0.01, \*\*\**P* < 0.001, \*\*\*\**P* < 0.0001 vs. corresponding control.

Since IL-6 is a glycol-protein whose secretion can be modulated by glycosylation^22^, we hypothesized that E-FOXP3 may regulate this post-translational modification. Co-immunoprecipitation assays revealed that E-FOXP3 overexpression specifically increased O-linked N-acetylgalactosamine (O-GalNAc)-modified IL-6, but not O-GlcNAc modifications (Fig. 5C). Among glycosyltransferases screened, GALNT1 and GALNT3 were transcriptionally upregulated by E-FOXP3 (Fig. 5D). To assess their functional roles, we overexpressed Flag-tagged IL-6 in hTERT-HPNE cells and knocked down GALNT1 or GALNT3. Although both knockdowns reduced global O-GalNAcylation, only GALNT1 depletion significantly diminished Flag–IL-6 O-GalNAc modification (Fig. 5E). Subsequent ChIP and luciferase reporter assays confirmed that E-FOXP3 directly binds the GALNT1, not GALNT3, promoter and transactivates its expression (Fig. 5F; Supplementary Fig. S6E, F). Critically, GALNT1 knockdown abrogated E-FOXP3-induced increases in intracellular O-GalNAcylated IL-6 and total O-GalNAc levels (Fig. 5G), and attenuated secretion of glycosylated IL-6 (Fig. 5H). Furthermore, GALNT1 silencing in hTERT-HPNE cells mitigated the pro-fibrotic activation of co-cultured PSCs (Fig. 5I). Together, these findings identify a non-canonical FOXP3-GALNT1-IL-6 axis wherein glycosylation serves as a key regulatory step in fibrosis initiation.

## Discussion

This study redefines the context-dependent role of FOXP3 beyond immune tolerance by establishing epithelial FOXP3 (E-FOXP3) as a pivotal initiator of pancreatic fibrocarcinogenesis through a non-immune mechanism. Unlike canonical drivers of fibrosis— such as immune cell infiltration or vascular inflammation^23,24^—E-FOXP3 operates quietly during early carcinogenesis, evading clinical detection. We demonstrate that E-FOXP3, though transiently expressed in rare epithelial subpopulations, functions as a critical trigger for fibrosis initiation during early neoplasia. Mechanistically, E-FOXP3 directly transactivates the glycosyltransferase GALNT1, catalyzing O-glycosylation of IL-6. This post-translational modification enables rapid secretion of glycosylated IL-6 (gIL-6), which then directly activates PSCs to induce inflammatory cancer-associated fibroblast (iCAF) formation (Fig. 6), unveiling a previously unrecognized upstream pathway that bridges epithelial dysfunction to stromal activation.

**Figure 6.**
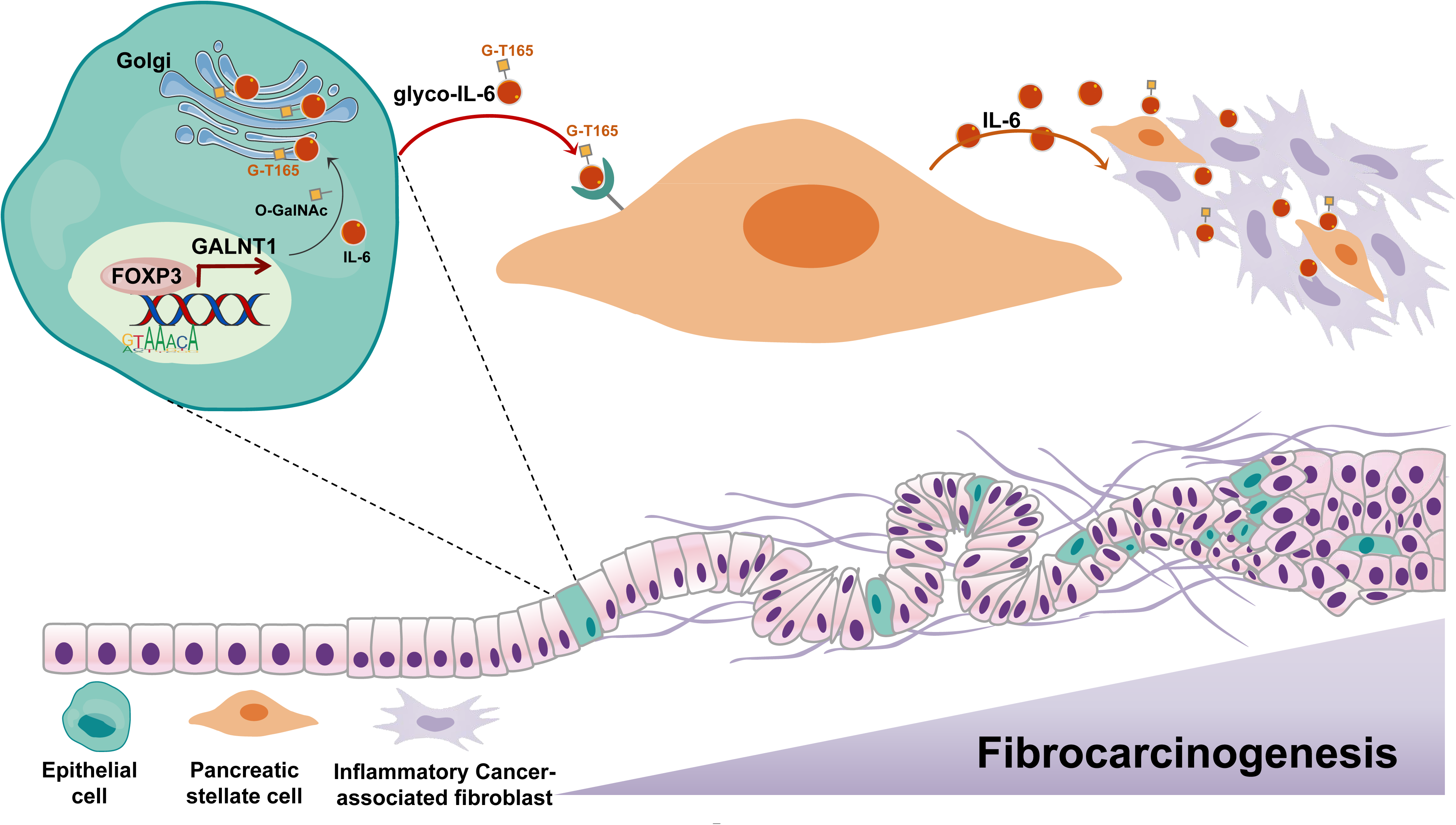
Model for the mechanism of epithelial FOXP3 drives pancreatic fibrosis via IL-6 glycosylation. Schematic cartoon depicting the mechanism for epithelial FOXP3 promotes stromal activation and fibrotic remodeling through IL-6 secretion.

Understanding the formation of iCAFs, which are key drivers of tumor progression, chemoresistance, and systemic effects, during early pancreatic preneoplasia is vital for future clinical strategies^25,26^. By revealing how epithelial FOXP3 drives inflammatory fibrosis during pancreatitis-to-cancer progression, this work provides a new mechanistic framework for early intervention. The identified pathway offers actionable therapeutic targets, while E-FOXP3 expression emerges as a promising biomarker for early detection. These insights collectively challenge the prevailing inflammation-centric view of pancreatic carcinogenesis^27–29^ and highlight epithelial-driven fibrosis as a targetable process in preneoplastic progression. Future efforts should focus on deciphering the upstream signals that reprogram epithelial cells toward this fibrogenic phenotype, which may unlock novel diagnostic and therapeutic strategies to intercept PDAC at its origin.

In conclusion, we unveil E-FOXP3 as master driver connecting epithelial dysfunction to stromal licensing in early pancreas pathobiology. By delineating a glycosylation-dependent signaling cascade that bypasses canonical immunity, our work provides a new mechanistic framework for understanding pancreatic cancer ’ s silent onset and aggressive evolution. Targeting this newly revealed axis offers a unified strategy to intercept fibrocarcinogenesis at its root.

## Acknowledgements

This work was supported by National Science Foundation of China (No. 82125026, 82330081 and 82472667), the Intergovernmental International Cooperation on Science and Technology Innovation Key Special Project of China (2024YFE0104100), the Natural Science Foundation of Shandong (No.ZR2025QC832). We thank Prof. Cai and colleagues from the Organ Transplantation Centre for their critical contributions to this study.

## Author Contributions

R.G., Z.Y. and H.R. conceived and designed the study and wrote the manuscript. R.G. and J.W. performed the experiments and data analysis. M.R. and W.Z. generated the genetically engineered mouse models and performed mIF staining. J.W. performed cell functional analysis. S.P. performed scRNA-seq analyses. H.S., J.C. and L.M. performed chromatin immunoprecipitation, co-immunoprecipitation assays and mass spectrometry analysis. Q.Y. and K.L. performed tissue collection. J.W. and J.X. assisted with histopathological evaluation. J.X. and M.L. provided materials. Z.Y., C.Z. performed cell culture. K.L., Q.Y., C.G., S.F. and T.Z. assisted with data interpretation and manuscript proofreading. S.Z. performed computational modeling and drug screening. A.S. provided critical linguistic editing. H.R. supervised this work. All authors reviewed and approved the final manuscript.

## Competing Interests

The authors declare no competing interests.

**Correspondence and requests for materials** should be addressed to He Ren.

## STAR METHODS

### KEY RESOURCES TABLE

| REAGENT or RESOURCE | SOURCE | IDENTIFIER |
| --- | --- | --- |
| Antibodies |  |  |
| FOXP3 | Abcam | ab215206 |
| CK19 | Abcam | ab52625 |
| α-SMA | Abcam | ab5694 |
| IL-6 | Abcam | AB290735 |
| IL-6 | CST | 12153S |
| IL-6 | Invitrogen | 701028 |
| ATAC2 | Abcam | ab5694 |
| GAPDH | Proteintech | 10494-1-AP |
| O-GalNAc | Sigma-Aldrich | SAB5202328-100UG |
| O-GlcNAc | Sigma-Aldrich | SAB5202310-100UG |
| GALNT1 | Proteintech | 21139-1-AP |
| GALNT3 | Proteintech | 16716-1-AP |
| FLAG | Proteintech | 20543-1-AP |
| PDGFRα+$\beta$ | Abcam | ab32570 |
| FAP | CST | 66562 |
| IL-6 Recombinant Rabbit<br>Monoclonal Antibody (4H16L21) | Invitrogen | 701028 |
| IL-6 (D3K2N) Rabbit mAb | CST | 12153S |
| Anti-alpha smooth muscle Actin<br>Antibody | abcam | ab5694 |
| GALNT1 Polyclonal antibody | Proteintech | 21139-1-AP |
| Biological samples |  |  |
| Primary tumour specimens and<br>muscle samples | Affiliated Hospital of<br>Qingdao University | N/A |
| Chemicals, peptide, and recombinant<br>proteins |  |  |
| Protease Inhibitor Cocktail | MCE | HY-K0011 |
| Stellate Cell Medium | ScienCell | 5301 |
| DMEM | Pricella | PM150210 |
| RPMI-1640 | Pricella | PM150110 |
| Foetal bovine serum | Gibco | A5669401 |
| Human Cytokine Array C1000 | RayBiotech | AAH-CYT-1000 |
| Human IL-6 ELISA Kit | Abcam | AB178013 |
| Siltuximab | Selleck | A2030 |
| Human IL-6 Recombinant Protein | PeproTech | 200-06-20UG |
| Mouse IL-6 Recombinant Protein | PeproTech | 216-16-10UG |
| eBioscience™ Protein Transport Inhibitor Cocktail | Thermo Fisher | 00-4980-93 |
| Human IL-6 Recombinant Protein, PeproTech® | thermofisher | 200-06 |
| SC144 | MCE | HY-15614 |
| Tocilizumab | Selleck | A2012 |
| Experimental models: Cell lines |  |  |
| hTERT-HPNE | ATCC | CRL-4023 |
| hPSCs |  | N/A |
| Deposited data |  |  |
| Critical commercial assays |  |  |
| BeyoChIP™ ChIP Assay Kit (Protein A/G Magnetic bead) | Beyotime | P2080S |
| BeyoMag™ Anti-Flag Magnetic Beads | Beyotime | P2115 |
| TruePrep Index Kit V2 for Illumina | Vazyme | TD202 |
| BeyoBlue™ Plus Coomassie Blue Super Fast Staining Solution | Beyotime | P0003M |
| ALphaTSA Multiplex IHC Kit For Tissue | Alphaxbio | AXT36100031 |
| SteadyPure Quick RNA Extraction Kit | AG | AG21023 |
| Evo M-MLV RT Kit with gDNA Clean for qPCR II | AG | AG11711 |
| SYBR Green Premix Pro Taq HS qPCR Kit | AG | AG11718 |

## Methods

### Experimental model and subject details

#### Cell lines

hTERT-HPNE, AsPC-1, PANC-1, and MIA-PaCa2 cell lines were purchased from the American Type Culture Collection (Manassas, VA, USA). Pancreatic stellate cells (PSCs) were kindly provided by Prof. Mingyang Liu (Cancer Hospital, Chinese Academy of Medical Sciences, Beijing). All cell lines underwent short tandem repeat (STR) authentication to confirm identity and ensure the absence of cross-contamination or misidentification. Mycoplasma contamination was ruled out using PCR-based testing. The hTERT-HPNE, PANC-1, and MIA PaCa-2 cell lines were maintained in Dulbecco’s modified Eagle medium (DMEM; Pricella, PM150210) supplemented with 10% foetal bovine serum (FBS; Gibco, A5669401) and 1% penicillin-streptomycin (Pricella, PB180120). AsPC-1 cells were cultivated in Roswell Park Memorial Institute (RPMI)-1640 medium (Pricella, PM150110). PSCs were cultured in a specialised stellate cell medium (ScienCell, 5301). All cells were maintained at 37 °C in a humidified atmosphere containing 5% CO_2_.

#### Animal models and experimental procedures

Transgenic mouse models, specifically RC and KFC, were established for this study. RC mice, conditionally expressing the *CAG-LSL-Foxp3-IRES-EGFP-Wpre-pA* transgene, were generated using CRISPR/Cas9 technology at Shanghai Model Organisms Center, Inc (Pudong District, Shanghai). Briefly, Cas9 mRNA and guide RNA (gRNA) were synthesised via *in vitro* transcription. A homologous recombination donor vector, containing a 3.3 kb 5′ and 3.3 kb 3′ homology arms, was constructed using In-Fusion cloning and microinjected into fertilised C57BL/6J mouse zygotes. Founder (F0) mice were screened by PCR, and positive animals were bred to produce F1 offspring. *Foxp3^fl/+^* and *Foxp3^fl/Y^* mice, generated on a C57BL/6 background, were engineered with loxP sites flanking exons 2–8 of the *Foxp3* gene. These mice were bred to obtain *Foxp3^flox/flox^* animals, which were subsequently crossed with KC mice to generate KFC mice. Control KC mice were maintained under identical conditions. All mice were monitored for general health, body weight (euthanasia was performed if weight loss exceeded 20% within 1 week), and overall survival. Animal procedures were approved by the Animal Ethics Committee of the Affiliated Hospital of Qingdao University and complied with institutional ethical standards (Approval number: AHQU-MAL20190510RH). Pancreatic tissues were collected for histological analysis at designated time points following euthanasia.

#### Clinical sample collection and ethics

Human pancreatic ductal adenocarcinoma (PDAC) tissue samples were gathered from patients at the Affiliated Hospital of Qingdao University. Samples were processed into formalin-fixed, paraffin-embedded (FFPE) blocks for subsequent analysis. All procedures involving human participants were conducted in accordance with Institutional Review Board (IRB) guidelines, with approval obtained under protocol number [QYFYWZLL29950, QYFYEC2023-151]. Informed consent was obtained from all participating patients prior to sample collection.

#### Stable cell line construction

Lentiviruses used to generate stable cell lines were purchased from OBiO Technology. Cells were infected with the lentiviral particles and subsequently selected using puromycin to isolate clones with successful viral integration. The selected cell populations were expanded, and stable FOXP3 expression was confirmed via quantitative PCR (qPCR) and western blot analysis

#### Co-culture assays

A total of 100,000 PSCs were seeded into each well of a 6-well plate and allowed to adhere overnight. The following day, the medium was replaced with 2.5 mL of fresh stellate cell medium. Subsequently, 200,000 HPNE cells were seeded into the upper chamber of a 0.4 µm transwell insert, which was then filled with 1.5 mL of medium. Both cell types were co-cultured for 24 h at 37 °C in a humidified incubator with 5% CO₂. After incubation, PSCs were harvested for immunofluorescence staining and qPCR analysis.

#### Immunocytochemistry

Cells were fixed using 4% paraformaldehyde and permeabilised using Triton X-100. Following a blocking step to reduce nonspecific binding, cells were incubated with the primary antibody overnight at 4 °C. After washing, cells were incubated with a fluorescently labelled secondary antibody (Alpha X Biotech, AXT36100031). For multiplex labelling, antibodies were washed out, and cells were re-blocked prior to incubation with additional primary antibodies as needed. Nuclei were counterstained using 4’,6-diamidino-2-phenylindole (DAPI), and slides were mounting for fluorescence microscopy and subsequent imaging.

#### ELISA

For ELISA, culture supernatants were collected and centrifuged to remove debris. Reagents were prepared according to the manufacturer’s instructions (Abcam, AB178013). In each detection well, 50 µL of each sample was added along with a cocktail of specific capture and detection antibodies targeting the protein of interest. Plates were incubated at 20–25 °C for 1 h. After washing, substrate solution was added, which reacted with the enzyme conjugated to the detection antibody, generating a colorimetric signal proportional to the concentration of target protein present. Finally, the reaction was stopped, and absorbance was measured at 450 nm using a microplate reader (Berthold, Tristar² S LB 942) to quantify the target protein in the supernatant.

#### Human cytokine array

After collecting culture supernatants from hTERT-HPNE cell, cytokine profiling was performed using the RayBio^®^ Human Cytokine Array C Series (RayBiotech, AAH-CYT-1000). Each array membrane was incubated with 1 mL of sample overnight at 4 °C, followed by detection of bound antigens according to the manufacturer’s instructions. The detection sensitivity of the antibodies on the array ranged 25–250,000 pg/mL. Detailed procedures are available in the user manual (https://doc.raybiotech.com/pdf/Manual/AAH-CYT-1000.pdf for details).

#### Quantitative reverse-transcription PCR (qRT-PCR)

Total RNA was extracted from cells using the RNA Quick Extraction Kit (AG, AG21023), and RNA concentration were measured prior to reverse transcription using a dedicated kit (AG, AG11711). The resulting cDNA was amplified with SYBR Green Master Mix (AG, AG11718) and gene-specific primers on a LightCycler^®^ 96 Instrument (Roche). glyceraldehyde-3-phosphate dehydrogenase (*GAPDH*) was used as a reference gene (forward primer: 5′-TGTGAGGGAGATGCTCAGTG-3′; reverse primer: 5′-TGTTCCTACCCCCAATGTGT-3′). Relative gene expression was calculated as fold change compared to control samples^30^. Thermal cycling conditions were as follows: initial denaturation at 95 °C for 30 s, followed by 40 cycles of denaturation at 95 °C for 5 s and annealing at 60 °C for 30 s.

#### Western blot analysis

Cell lysates were prepared using radioimmunoprecipitation assay lysis buffer supplemented with protease inhibitors as previously described^31^. Protein concentrations were determined using the bicinchoninic acid (BCA) Protein Assay Kit (Solarbio, Cat# PC0020). Equal amounts of protein were separated on 15% SDS-PAGE gels and transferred to a polyvinylidene fluoride membrane (Millipore, Cat# IPVH00010). Membranes were blocked with 5% non-fat milk in TBS-T (0.1% Tween-20 in Tris-buffered saline) for 2 h at 25 °C, then incubated overnight at 4 °C with primary antibodies against alpha smooth muscle actin (α-SMA, 1:1000), IL-6 (1:1000), O-GlcNAc (1:1000), GAPDH (1:5000), and FOXP3 (1:1000). After washing, membranes were incubated with horseradish peroxidase-conjugated secondary antibodies (1:10,000) for 2 h at 25 °C. Protein bands were visualised using enhanced chemiluminescence (ECL) reagents, following the manufacturer’s protocol.

#### Chromatin immunoprecipitation assay (ChIP)

ChIP was performed in hTERT-HPNE cells using an anti-FOXP3 antibody and the Magnetic Chromatin Immunoprecipitation System (Life Technologies), following the manufacturer’s instructions. Following immunoprecipitation and antibody removal, target DNA fragments were amplified by PCR with following thermal cycling conditions: initial denaturation at 95 °C for 30 s, followed by 40 cycles of denaturation at 95 °C for 5 s and annealing at 60 °C for 30 s. The following forward and reverse primer pairs were used to amplify FOXP3-binding sites and p21 control site: Site1 (5′-TGCTTGCAACAAGTTCTTTGC-3′ and 5′-ACAGTTGTGAAAAGTGCATTTGA-3′); Site2 (5′-TTTTCAGCTGCATATAATCTTATGGGA-3′ and 5′-CATACAGTGGACTACTACTCAGTAA-3′); Site3 (5′-ACCTGTACAGCATGGGACTG-3′ and 5′-TGTCGTTGGGCAATGGGTTA-3′); Site4 (5′-ACGGTAGAATTATGTGCAAACAGC-3′ and 5′-AGCGCAGGAAAATAGTACCTT-3′); Site5 (5′-TTTTTCGGACAGCCTTTTGGT-3′ and 5′-AGACAAAACCCTGGCTCTGT-3′); and P21-site (positive control) (5′-AGTCAGTTCCTTGTGGAGCC-3′ and 5′-GGACACGCAGGGACACAC-3′).

#### Co-immunoprecipitation (Co-IP) assay

Co-IP was performed using the Pierce™ Classic Magnetic IP/Co-IP Kit (Beyotime, #P2115), as previously described^32^. Briefly, cell lysates were incubated with pre-washed Pierce anti-FLAG protein A/G magnetic beads for 1 h at room temperature. After incubation, antigen–antibody–bead complexes were collected using a magnetic stand and washed thoroughly to remove non-specific binding. The bound proteins were eluted and analysed by western blotting.

#### Single-cell RNA sequencing (scRNA-seq) data analysis

Mouse pancreatic tissues were minced and enzymatically digested in a solution containing collagenase IV (2 mg/mL; Sigma-Aldrich) and deoxyribonuclease I (1 mg/mL; Sigma-Aldrich) at 37 °C for 30 min. The resulting cell suspension was mixed with 2% FBS in phosphate-buffered saline (PBS) and centrifuged at 300 *g* for 5 min at 4 °C. The pellet was then treated with red blood cell lysis buffer (BD) for 3 min at room temperature, followed by centrifugation at 300 *g* for 5 min at 4 ℃. The cells were resuspended in PBS and loaded onto a 10X Genomics GemCode Single-cell instrument to generate single-cell gel bead-in-emulsion (GEMs). GEM generation, barcoding, cDNA synthesis, and library construction were performed using Chromium Next GEM Single Cell 3′ Reagent Kits v3.1 according to the manufacturer’s protocol. Libraries were sequenced on an Illumina Novaseq 6000 platform (Gene Denovo Biotechnology Co., Guangzhou, China). Raw base call (BCL) files were converted to FASTQ files, aligned, and quantified using Cell Ranger software (version 3.1.0, 10X Genomics). The resulting raw gene expression matrices were processed using the Seurat R package (v4.4.1). Cells were filtered based on the following criteria: unique molecular identifiers (UMI) between 200 and 25,000, gene count (nGene) between 200 and 6,000, and mitochondrial gene content < 5%. The data were normalised to a total of 10,000 counts per cell. Cell doublets were removed using the DoubletFinder R package (version 2.0.3). Gene expression was normalised using the ‘LogNormalize’ method, which normalises each gene’s expression, multiplies this by a scale factor (default 10,000), and log-transforms the result. To minimise the effects of batch effects and behavioural conditions on clustering, we used Harmony, an algorithm that projects cells into a shared embedding in which cells are grouped by cell type rather than dataset-specific conditions, to aggregate all samples^33^. The Harmony algorithm inputs a PCA embedding of cells along with their batch assignments and returns the batch-corrected embedding. The number of principal components (PCs) was adjusted to 30 to generate cell clusters, which were then visualised using Uniform Manifold Approximation and Projection (UMAP).

#### Cell type annotation and differentially expressed gene (DEG) analysis

Cell types were annotated based on canonical marker gene expression. The ‘FindAllMarker’ function in the Seurat package was used to identify DEGs for each cell cluster using default parameters. Clusters with similar gene expression signatures were grouped and annotated as the same cell type. For targeted comparisons between specific clusters, the ‘FindMarkers’ function was employed to identify DEGs.

#### Haematoxylin and eosin (H&E) staining

Pancreatic tissues were harvested, fixed in 10% formalin for 2 days, followed by paraffin embedding. Tissue sections were deparaffinised in xylene followed by rehydration through a graded ethanol series. Nuclei were stained with haematoxylin, followed by rinsing and differentiation in hydrochloric acid-ethanol. Cytoplasmic components were counterstained with eosin. Sections were dehydrated, cleared in xylene, and mounted with coverslips. Lesion areas and histological grades were independently evaluated by two senior pathologists. A third pathologist was consulted in cases of discordance to reach a consensus.

#### Multiplex immunofluorescence (mIF)

Paraffin-embedded tissue sections were processed using the AlphaTSA Automated Multiplex IHC Kit. Following deparaffinisation and rehydration, antigen retrieval was performed by heat-induced epitope retrieval using citrate or ethylenediaminetetraacetic acid solution and microwave heating. After blocking nonspecific binding sites, sections were incubated overnight at 4 °C in a humidified chamber with primary antibodies. After washing, the sections were incubated with secondary antibody and fluorescent dyes sequentially. Fluorescent images were acquired using a Zeiss fluorescence microscope. Quantitative analysis of multiplex signals was performed using HALO AI™ image analysis software.

#### Immunohistochemistry (IHC)

Formalin-fixed, paraffin-embedded tissue sections were deparaffinised and rehydrated, followed by antigen retrieval (as described above) at 95 °C for 15 min. After quenching endogenous peroxidase activity and blocking nonspecific binding sites, sections were incubated overnight with primary antibodies at 4 °C. After washing, sections were then incubated with biotinylated secondary antibodies, followed by detection using 3,3’-diaminobenzidine (DAB) substrate. Haematoxylin was used for counterstaining, followed by dehydration, xylene clearing, and mounting with coverslips.

#### Alcian Blue staining

Deparaffinised and rehydrated sections were treated with 0.1 M acetic acid, followed by staining with Alcian Blue solution. After thorough washing, nuclei were counterstained red. The sections were then dehydrated and mounted with coverslips.

#### Sirius Red staining

Tissue sections were deparaffinised and rehydrated, then stained with haematoxylin to label nuclei. After rinsing, sections were stained with Sirius Red to visualise collagen fibres. Sections were then washed, dehydrated, and mounted with coverslips.

#### Masson’s Trichrome staining

Deparaffinised and rehydrated sections were stained with haematoxylin overnight and incubated at 65 °C. After washing and differentiation in hydrochloric acid alcohol, the sections were sequentially stained with phosphomolybdic acid and aniline blue. Following dehydration, coverslips were mounted.

#### Statistical analyses and reproducibility

Statistical analyses were performed as described in the corresponding figure legends. A *P* value less than 0.05 was considered statistically significant. Sample sizes were not predetermined using statistical methods. For details regarding randomisation and blinding, refer to the Reporting Summary. Most experiments were not randomised, as they did not involve interventions requiring random assignment. Unless otherwise indicated, statistical analyses were performed using GraphPad Prism software (v8.0.1).

