## Supplemental Figure legends for "Epithelial FOXP3 Drives Pancreatic Fibrosis through O-glycosylated IL-6"

**Supplementary Figures 1-8**

**Figure S1. Phenotypic and histological comparison of KC and KFC mice.**

**Figure S2. Ablation of FOXP3$^+$ epithelial cells attenuates pancreatic fibrosis.**

**Figure S3. Phenotypic and genotypic characterisation of R and RC mic.**

**Figure S4. Identification and spatial localization of CK19$^+$FOXP3$^+$ and IL-6$^+$ cells in pancreatic tissues.**

**Figure S5. IL-6 mediates PSC activation *in vitro*.**

**Figure S6. FOXP3 promotes IL-6 secretion through transcriptional regulation of GALNT1 without affecting IL-6 expression.**

**Figure S1: Phenotypic and histological comparison of KC and KFC mice. A and B,** H&E staining (A) and FOXP3 IHC staining (B) of human PanIN tissues from donor samples. Scale bars: 50 μm. **C**, Schematic of the experimental design. **d-g,** FOXP3 IHC staining of various mouse tissue samples: pancreatic tissue from WT mice **(**D**),** chronic pancreatitis tissue from WT mice (E), pancreatic tissue from *Kras$^{LSL\text{-}G12D/+}$*; *Pdx1*-Cre (KC) mice (F), and chronic pancreatitis tissue from KC mice (G). WT, wild-type; KC, *Kras$^{LSL\text{-}G12D/+}$; Pdx1*-Cre mice. Scale bars: 50 μm. **H**, Cytoplasmic FOXP3$^+$ epithelial cell counts in the pancreases of mice. **I**, Nuclear FOXP3$^+$ epithelial cell counts in the pancreases of mice.

**Figure S2: Ablation of FOXP3$^+$ epithelial cells attenuates pancreatic fibrosis. A,** Schematic representation of KC and KFC mice. **B,** Longitudinal analysis of body weight changes in KC and KFC mice. KC, *LSL-Kras*$^{G12D}$; *Pdx1-Cre* mice; KFC, *Kras*$^{G12D}$*; FOXP3*$^{flox/flox}$*; Pdx1-Cre* mice. Two-way ANOVA was performed to compare the body weight; *, $P = 0.035$. **C,** Pancreatic weight-to-body weight ratio in KC versus KFC mice (unpaired two-tailed *t*-tests). **D,** Gross morphological comparison of the pancreas between KC and KFC mice. **E**, Histological evaluation of pancreatic lesions in age-matched KC and KFC mice ($n = 8$) based on H&E-stained sections and categorised as normal pancreas (Nor), PanIN1, PanIN2, and PanIN3. **F,** Alcian Blue staining of caerulein-induced chronic pancreatitis in KC and KFC mice. Scale bar: 50 μm. **G,** Bar graphs showing the percentage of Alcian Blue–positive lesions in the pancreases of mice ($n = 8$). **H,** Sirius Red staining of caerulein-induced chronic pancreatitis in KC and KFC mice. Scale bar: 50 μm. **I,** Percentage of Sirius Red–positive lesions in the pancreases of mice ($n = 8$). **J,** Masson's trichrome staining of caerulein-induced chronic pancreatitis in KC and KFC mice. Scale bar: 50 μm. **K,** Percentage of Masson's trichrome–positive staining in the pancreases of mice ($n = 8$). **L,** Bar graphs showing the percentage of α-SMA$^+$ cells relative to PDGFRβ$^+$ cells in the pancreases of mice ($n = 8$).

**Figure S3: Phenotypic and genotypic characterisation of R and RC mic. A,** Gross morphological appearance of the pancreas in R and RC mice. **B,** Longitudinal body weight changes in R and RC mice. Two-way ANOVA was performed to compare the body weight; ****, $P < 0.0001$. **C,** Pancreatic weight-to-body weight ratio in R versus RC mice (unpaired two-tailed *t*-tests). **D,** Representative videos illustrating the external appearance and activity levels of R and RC mice. Data in (D) are represented as mean ± SD for n = 3 biological replicates. ****$P < 0.0001$ determined by a paired two-tailed *t* test. **E** and **F,** Histological grading of pancreatic intraepithelial neoplasia (PanIN) lesions at stages PanIN1, PanIN2, and PanIN3 in pancreatic tissues from RC mice. Scale bar: 50 μm. Data in (F) is represented by one-way ANOVA with Tukey' s post-hoc test. ****$P < 0.0001$ vs. corresponding control.

**Figure S4: Identification and spatial localization of CK19$^+$FOXP3$^+$ and IL-6$^+$**

**cells in pancreatic tissues. A,** Heat map showing cell type–enriched genes. Each column represents an individual cell, and each row represents a cell type–specific signature gene. The colour scale (red to blue) indicates relative gene expression levels (high to low). **B**, Spatial plot showing the distribution of $CK19^{+}FOXP3^{+}$ cells and $IL\text{-}6^{+}$ cells in mouse pancreatic tissues. Scale bar: 100 μm. **C**, Spatial plot showing the distribution of $CK19^{+}FOXP3^{+}$ cells and $IL\text{-}6^{+}$ cells in human PDAC pancreatic tissues. Scale bar: 100 μm.

**Figure S5: IL-6 mediates PSC activation *in vitro*. A,** Quantitative analysis of the fluorescence intensity of PSC activation markers after 24 h of co-culture. **B**, ELISA validation of IL-6 levels based on cytokine array results. **C**, Western blot validation of IL-6 and α-SMA levels based on cytokine array results. **D**, Relative mRNA levels of PSC activation markers after 24 h of IL-6 stimulation at different concentrations. **E**, Time-course analysis of mRNA levels of PSC activation markers after treatment with 100 ng/mL IL-6. Data in (A) and (B) are represented by a paired two-tailed *t* test. (D) and (E) are represented by one-way ANOVA with Tukey’s post-hoc test. *$P < 0.05$, **$P < 0.01$, ***$P < 0.001$, ****$P < 0.0001$ vs. corresponding control.

**Figure S6: FOXP3 promotes IL-6 secretion through transcriptional regulation of GALNT1 without affecting IL-6 expression. A,** RT–qPCR analysis of IL-6 mRNA in hTERT-HPNE cells infected with control lentivirus (LV-Con) or FOXP3-overexpressing lentivirus (LV-FOXP3). **B,** Chromatin immunoprecipitation (ChIP) assay using anti-FOXP3 antibody to assess binding to the IL-6 promoter in hTERT-HPNE cells. **C,** IL-6 mRNA stability measured by RT–qPCR in LV-Con- or LV-FOXP3-infected hTERT-HPNE cells after treatment with actinomycin D at indicated time points. **D,** Quantification of intracellular and extracellular IL-6 protein levels by immunoblotting. **E,** ChIP assay evaluating FOXP3 binding to the GALNT3 promoter. **F,** Dual-luciferase reporter assay of GALNT1 promoter activity in the presence or absence of FOXP3. Data are presented as mean ± SD; n = 3 biological replicates. *$P < 0.05$, **$P < 0.01$, ***$P < 0.001$; paired two-tailed t-test.
