## Supplementary figures and images for "Epithelial FOXP3 Drives Pancreatic Fibrosis through O-glycosylated IL-6"

### Supplemental Figures

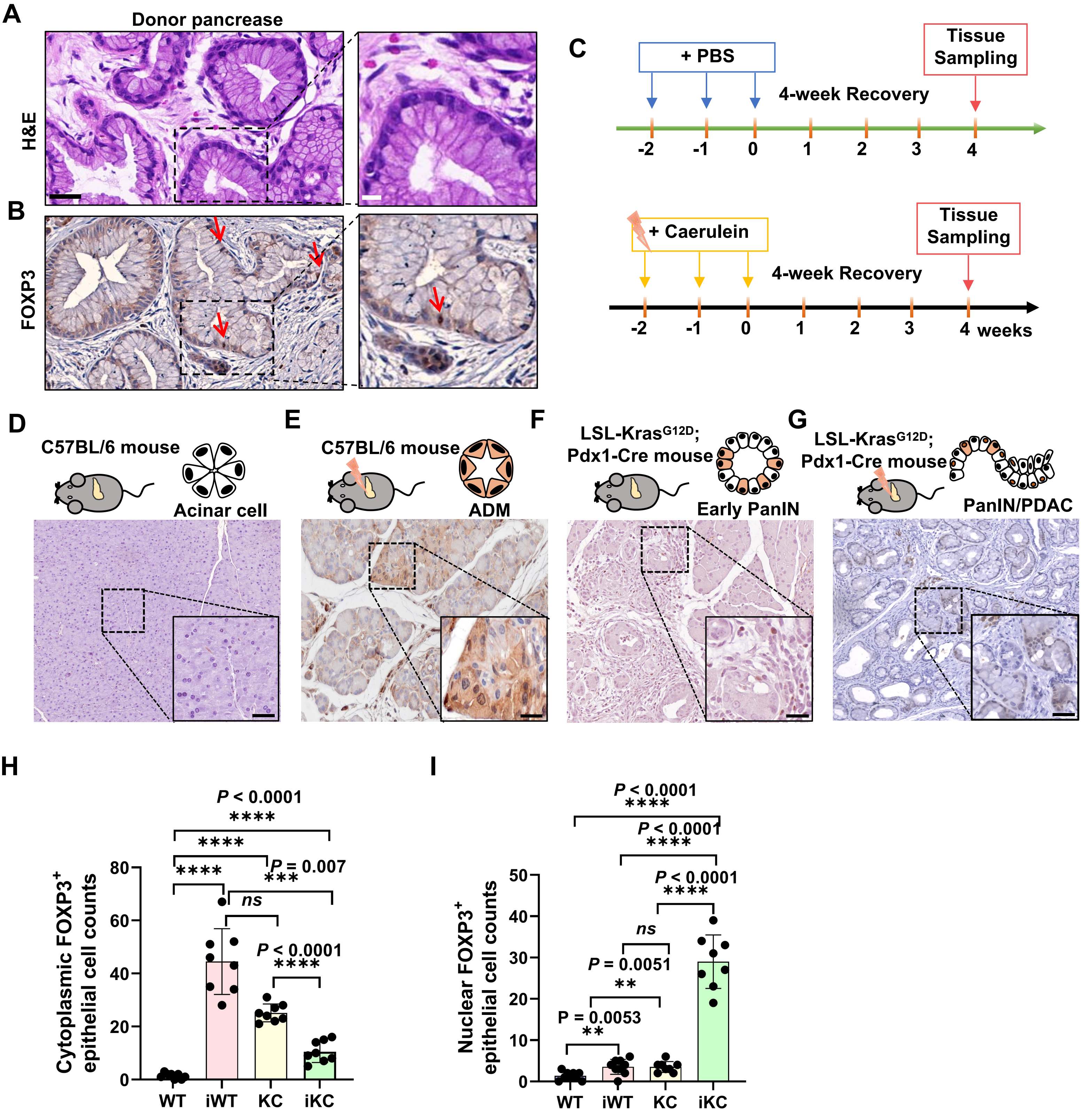

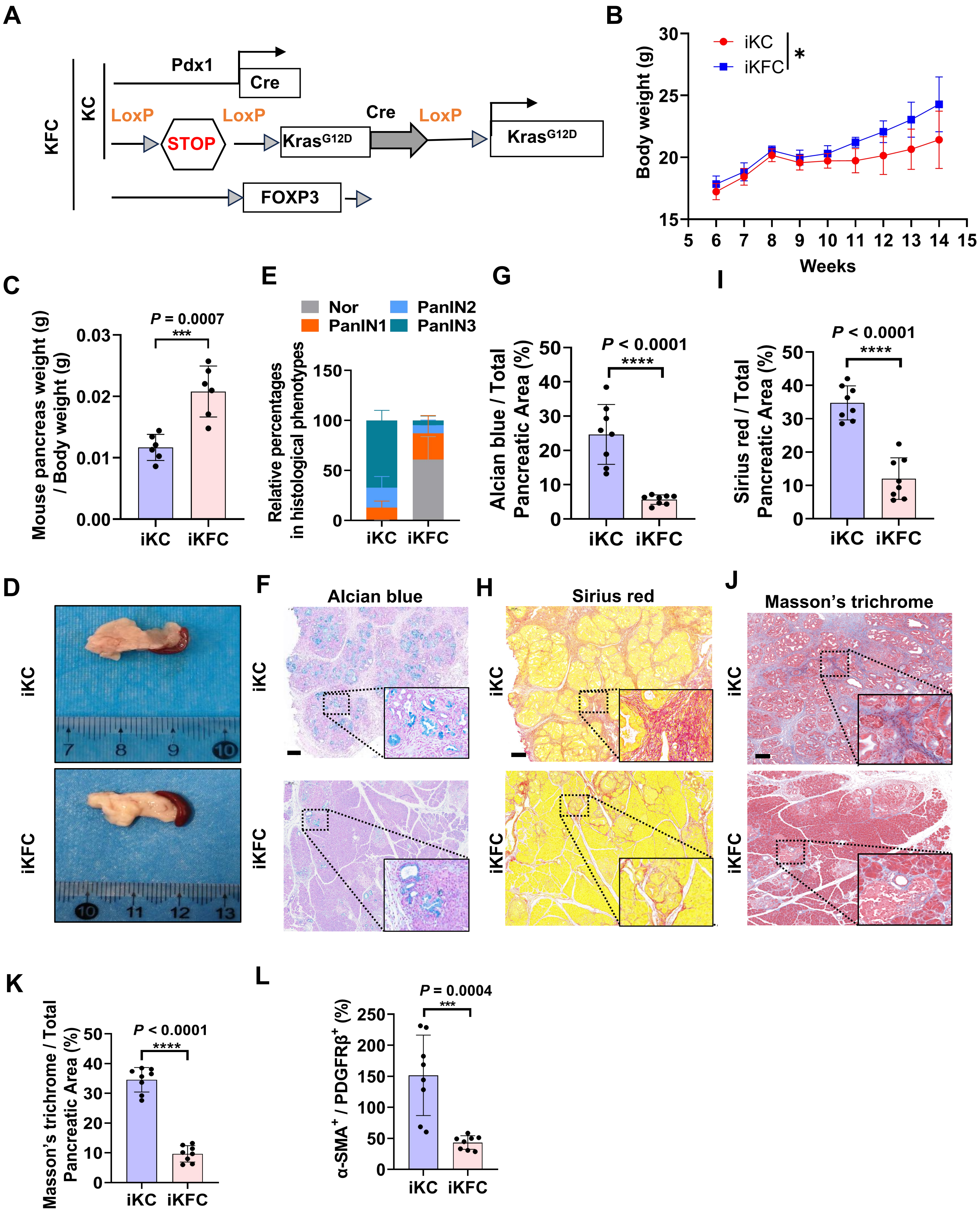

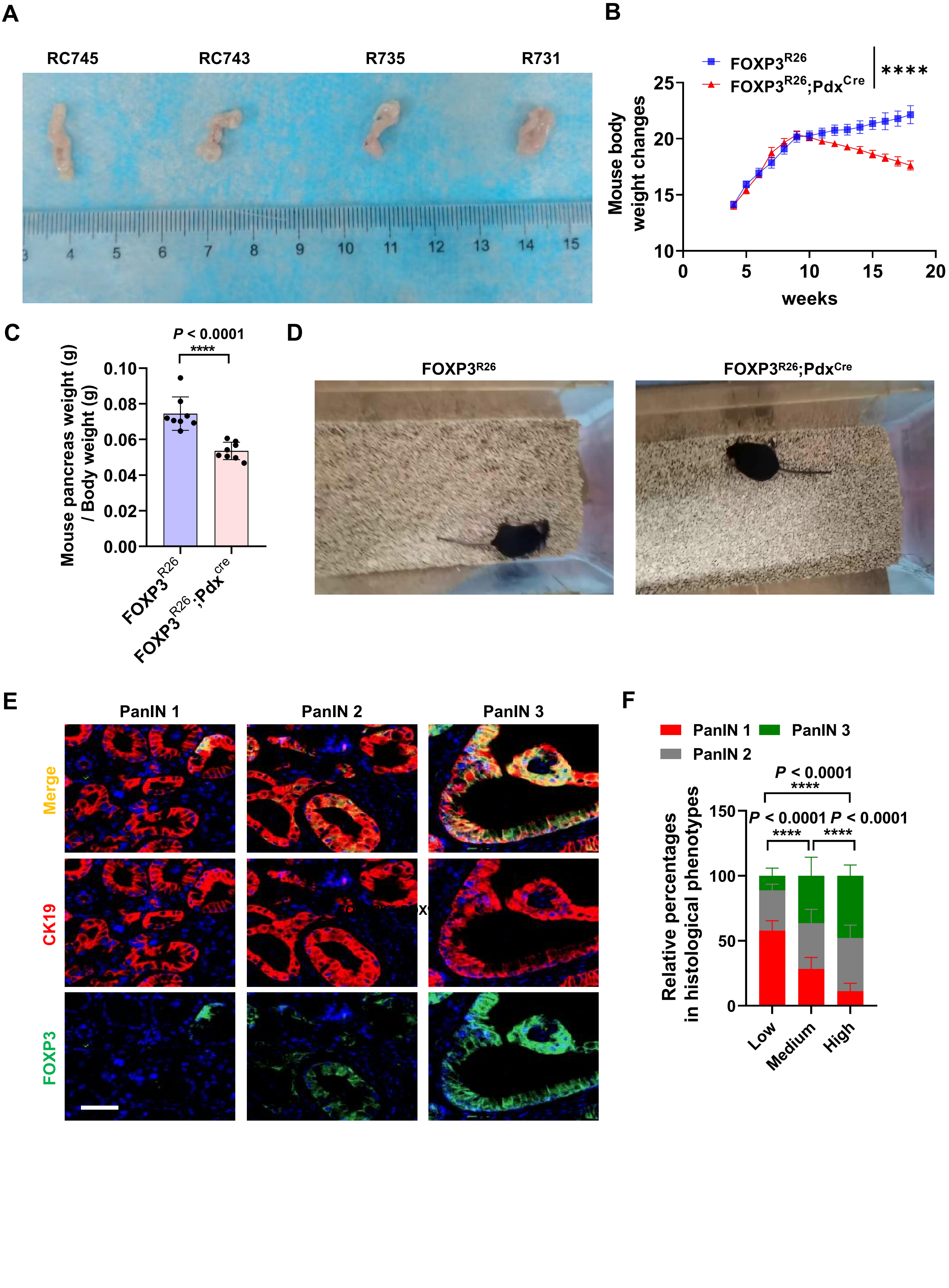

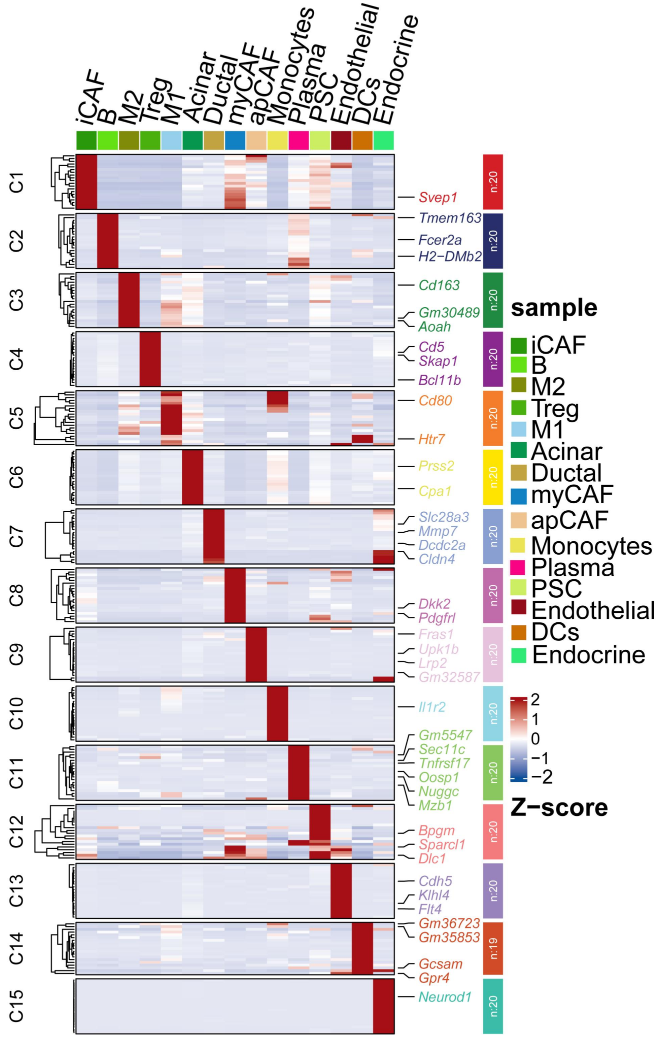

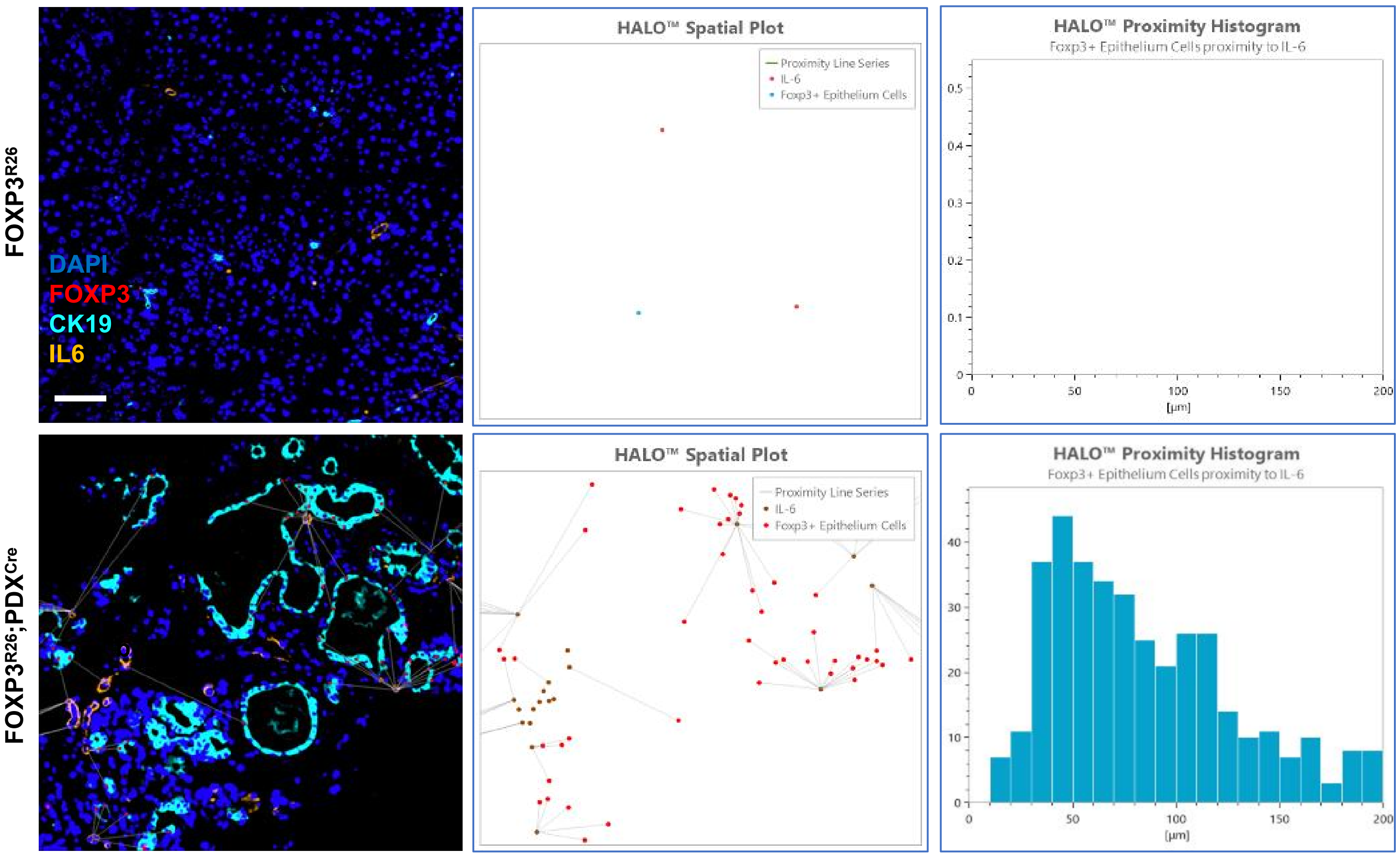

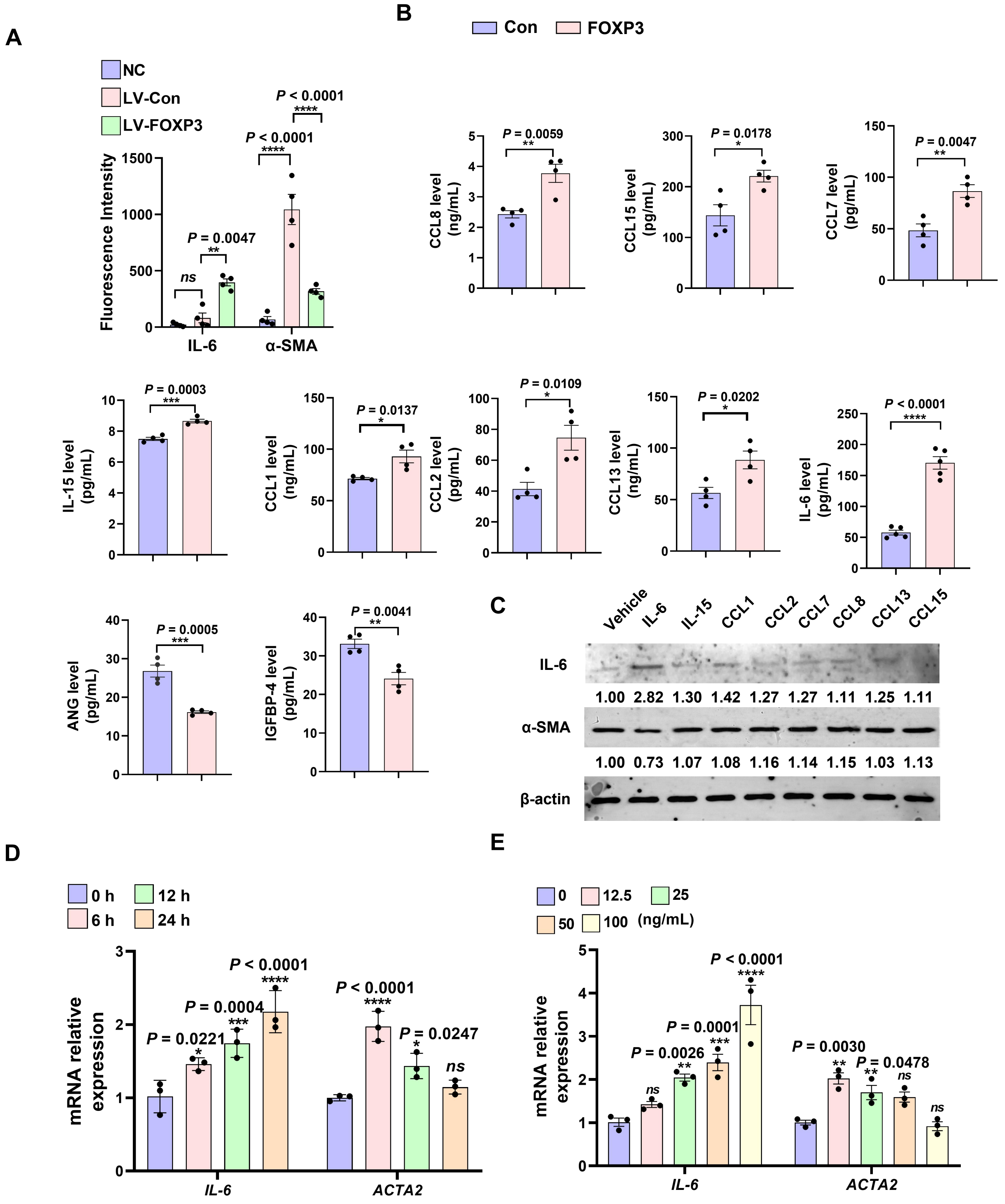

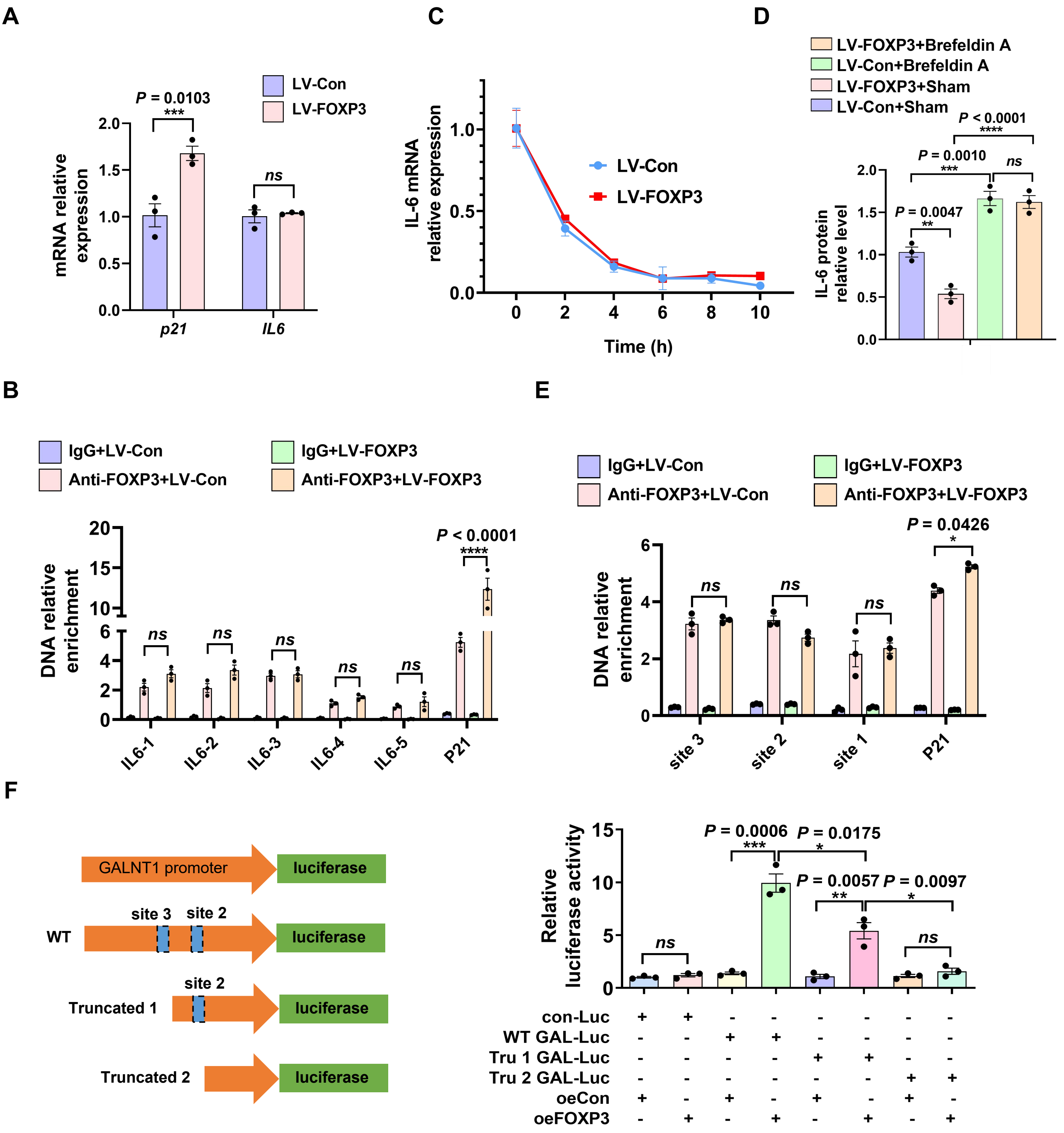
